# Climate, host and geography shape insect and fungal communities of trees

**DOI:** 10.1101/2023.04.17.537195

**Authors:** Iva Franić, Eric Allan, Simone Prospero, Kalev Adamson, Fabio Attorre, Marie Anne Auger-Rozenberg, Sylvie Augustin, Dimitrios Avtzis, Wim Baert, Marek Barta, Kenneth Bauters, Amani Bellahirech, Piotr Boroń, Helena Bragança, Tereza Brestovanská, May Bente Brurberg, Treena Burgess, Daiva Burokienė, Michelle Cleary, Juan Corley, David R Coyle, György Csóka, Karel Černý, Kateryna Davydenko, Maarten de Groot, Julio Javier Diez, H. Tuğba Doğmuş Lehtijärvi, Rein Drenkhan, Jacqueline Edwards, Mohammed Elsafy, Csaba Béla Eötvös, Roman Falko, Jianting Fan, Nina Feddern, Ágnes Fürjes-Mikó, Martin M. Gossner, Bartłomiej Grad, Martin Hartmann, Ludmila Havrdova, Miriam Kádasi Horáková, Markéta Hrabětová, Mathias Just Justesen, Magdalena Kacprzyk, Marc Kenis, Natalia Kirichenko, Marta Kovač, Volodymyr Kramarets, Nikola Lacković, Maria Victoria Lantschner, Jelena Lazarević, Marianna Leskiv, Hongmei Li, Corrie Lynne Madsen, Chris Malumphy, Dinka Matošević, Iryna Matsiakh, Tom W. May, Johan Meffert, Duccio Migliorini, Christo Nikolov, Richard O’Hanlon, Funda Oskay, Trudy Paap, Taras Parpan, Barbara Piškur, Hans Peter Ravn, John Richard, Anne Ronse, Alain Roques, Beat Ruffner, Alberto Santini, Karolis Sivickis, Carolina Soliani, Venche Talgø, Maria Tomoshevich, Anne Uimari, Michael Ulyshen, Anna Maria Vettraino, Caterina Villari, Yongjun Wang, Johanna Witzell, Milica Zlatković, René Eschen

**Author notes:** corresponding author: Iva Franić, Current address: Southern Swedish Forest Research Centre, Swedish University of Agricultural Sciences, Alnarp, SE.

## Abstract

Non-native pests, climate change, and their interactions are likely to alter relationships between trees and tree-associated organisms with consequences for forest health. To understand and predict such changes, factors structuring tree-associated communities need to be determined. Here, we analysed the data consisting of records of insects and fungi collected from dormant twigs from 155 tree species at 51 botanical gardens or arboreta in 32 countries. Generalized dissimilarity models revealed similar relative importance of studied climatic, host-related and geographic factors on differences in tree-associated communities. Mean annual temperature, phylogenetic distance between hosts and geographic distance between locations were the major drivers of dissimilarities. The increasing importance of high temperatures on differences in studied communities indicate that climate change could affect tree-associated organisms directly and indirectly through host range shifts. Insect and fungal communities were more similar between closely related vs. distant hosts suggesting that host range shifts may facilitate the emergence of new pests. Moreover, dissimilarities among tree-associated communities increased with geographic distance suggesting that human-mediated transport may facilitate the introductions of new pests. The results of this study highlight the need to limit the establishment of tree pests and increase the resilience of forest ecosystems to changes in climate.

## Introduction

Trees support a huge diversity of antagonistic and mutualistic organisms^1^, including numerous insects and fungi. Of these, herbivorous insects and plant pathogenic fungi can reduce tree growth and cause large-scale mortality^2^, while certain fungal mutualists increase the resistance of trees to abiotic and biotic stresses^3,4^. Many tree-associated fungi are also known as saprotrophs which have an important role in terrestrial carbon and nutrient cycling^5^. Interactions between trees and tree-associated organisms are likely to be strongly affected by the on-going global change (i.e., invasions by non-native pests and climate change) and changes in these interactions would likely have detrimental consequences for forest ecosystems. However, the extent of the expected effects and their consequences are unknown. In recent years, the number of non-native tree pests (mostly herbivorous insects and plant pathogenic fungi) has been increasing dramatically^6,7^ with serious consequences for tree and forest health^8,9^, and it is likely that this trend will continue in the future^10^. Furthermore, climate change could facilitate the range expansion of tree pests, and it might also create stressful conditions that make host plants more susceptible to antagonists, resulting in more pest outbreaks^11–13^. In order to understand and mitigate the consequences of global change on trees and forests, we need to understand the large-scale drivers of diversity of tree-associated organisms.

Large-scale drivers of variation in organism communities can be studied by focusing on mechanisms that drive differences in species composition and abundance between sites (i.e., β-diversity). The most important mechanisms shaping species assemblages, and variations between them are: 1) matching between the abiotic environment and the organism, 2) dispersal limitations which are linked to organisms’ ability to spread and to geographic barriers^14,15^, and 3) interactions with living components of the environment^16^. For example, in the case of tree-associated insects and fungi, specific climatic conditions might select for well adapted species that can tolerate present conditions, but climate might also determine the surrounding vegetation which will have an effect on tree-associated organisms. Host imposed filtering is based on the degree of matching between host traits, such as biochemical and physical defenses and organism’s ability to overcome them. Also, as a consequence of large geographic distances and geographic barriers between locations, tree-associated organisms might be spatially structured. Understanding the relative importance of climatic, host-related and geographic factors for shaping the communities of tree-associated organisms on large scales is crucial for predicting the impact that global change will have on these communities.

Determining the relative contribution of climatic variables in shaping the β-diversity of tree-associated communities would allow predictions about how climate change will alter them. Many insects and fungi show strong physiological adaptations for specific climatic conditions and their diversity and community composition could therefore directly respond to climate change. Community composition of tree-associated fungi reflects both temperature seasonality and climate (mean annual temperature and precipitation) in tropical forests^17^. Similarly, insect communities were found to be significantly affected by climatic factors, with gradients in temperature parameters being more important in driving β-diversity than gradients in precipitation parameters^18^. However, significant, but low impact of climatic variables on community composition of known tree pests was shown in a recent global analysis of tree pests^19^. Besides the direct effects, climate change could also indirectly affect associated communities by driving range shifts in tree species which is why it is also important to quantify the importance of host-related variables in shaping these communities.

Different tree species will harbor different associated taxa and turnover in fungal and insect communities might be predicted by differences in host traits^20,21^ or by phylogenetic relatedness^19^, which reflects the adaptation of these organisms to their hosts during long periods of co-evolution^21,22^. A phylogenetic signal in host association, i.e., a higher community similarity between closely related hosts, was previously shown for tree fungi^23–25^ as well as for herbivorous insects^26^. We might predict that antagonistic taxa are more specialized and show a stronger response to phylogeny than many mutualists^24^ or decomposers. In contrast, wood decay fungi might respond strongly to key functional traits such as wood density^27^. Wood density in fact could be an important factor for all tree-associated organisms as it is a major trait differentiating trees with different ecological strategies^28^. Also, hemisphere of origin of the tree species might reflect evolutionary adaptations of a tree species to the environment in which it evolved^29^, and thus could be an important factor structuring communities of insect and fungi. Furthermore, fungal communities associated with living trees are often dominated by only a few taxa^22^ and these dominant fungi may respond more strongly to host phylogeny and traits compared to the rarer taxa. In natural environments, the various drivers of insect and fungal communities are often confounded, as tree species composition is determined by abiotic conditions and dispersal limitation. Sampling of many host tree species across a wide geographic range, and in contrasting environmental conditions, is thus crucial for assessing the factors that shape β-diversity of tree-associated insects and fungi.

At large spatial scales, dispersal limitation might also structure tree-associated communities. For example, distinct insect and fungal communities might appear in association with trees growing in the Northern vs. Southern hemisphere^30^ and this might be because of the reasons such as geographic barriers or wind currents not crossing the equator, with both possibly limiting the exchange of species between the hemispheres. Similarly, intercontinental differences in insect and fungal communities might exist due to geographic barriers among continents which limit the spread of these organisms^31^. In the past centuries humans have reduced dispersal barriers by moving species, for example by growing trees outside their native ranges. Non-native trees may harbour fewer herbivorous insect species and plant pathogenic fungi than native trees, as the non-natives leave many of their specialists behind when they are introduced^32,33^. The degree of geographic structure in the community composition of tree-associated organisms is poorly known but is important to predict how human-mediated global exchange of trees might further spread insects and fungi around the globe.

The vast majority of studies using biological sampling of tree-associated insects and fungi from multiple tree species, which is necessary to generalize across hosts, determined the relative importance of different factors on their community assemblages by focusing on small geographic or environmental scales. For example, drivers of community composition of tree-associated insects and fungi were studied for multiple hosts in tropical^17,24,34,35^, boreal^36^ or temperate^37,38^ regions, but rarely in multiple climatic regions simultaneously^39^. Furthermore, drivers of community composition were rarely assessed for insects and fungi collected in a standardized way from the same samples^37^. However, to be able to understand and predict the impacts of global change the assessment needs to be done for both insects and fungi, on a large geographic scale across climatic regions, and for many hosts simultaneously, as large-scale processes determining the assemblages might be different than ones on the local scales, and might vary across organism groups.

We studied herbivorous insects and fungi in dormant twig samples (pooled wood, buds and needles/evergreen leaves from 20 twigs) from 155 angiosperm and gymnosperm tree species collected in botanical gardens or arboreta in 32 countries across the globe (Fig. 1). The dataset revealed the diversity of tree-associated herbivorous insects and fungi across broad geographic and climatic gradients and for many host taxa^40^ which allowed us to assess the relative impact of local climatic variables (i.e., mean annual temperature, mean annual precipitation, temperature seasonality), host-related variables (i.e., host phylogeny, wood density, hemisphere of origin, native/non-native status of a tree species (“native vs. non-native range”)) and geographic variables (i.e., geographic distance between sampling locations, hemisphere of collection) on β-diversity of herbivorous insects and fungi which might be moved through trade of plant material^40^. To target the organisms that might be associated with plant movements, dormant tree twigs were sampled in winter months because woody plants are often traded in winter, and mainly as budwood (i.e., twigs without leaves and roots attached). We used generalized dissimilarity models (GDMs) to analyse incidence-based and abundance-weighted β-diversity of herbivorous insects and all, saprotrophic, symbiotrophic and plant pathogenic fungi. GDMs allowed us to estimate the unique effect of each variable on β-diversity and to test for non-linear relationships. In order to test whether rare and common species differed in their responses, we calculated several measures of β-diversity which give different weightings to species abundance, based on Hill numbers^41^.

**Figure 1.**
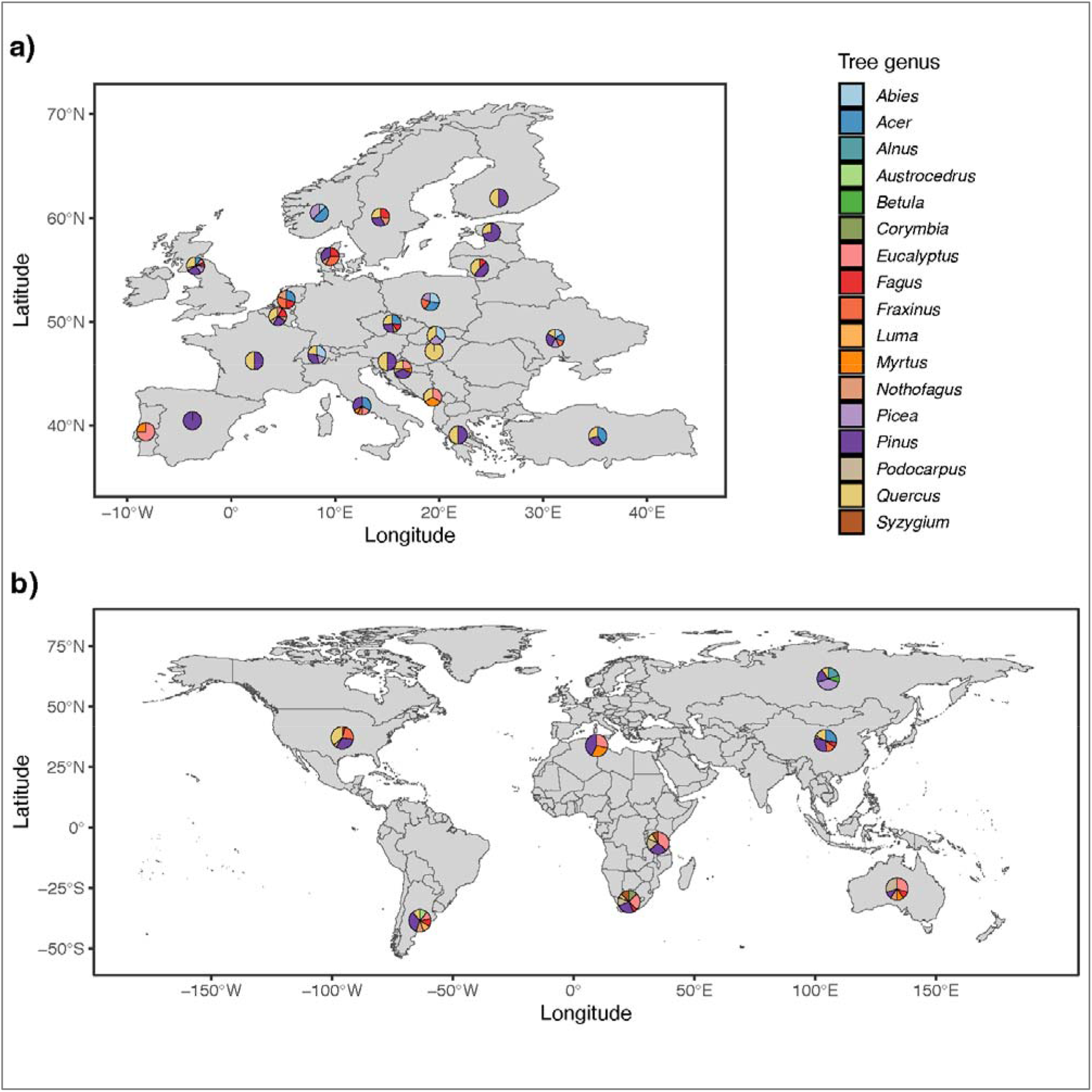
Geographic distribution and host species information for study samples across countries. Maps are shown for countries within Europe (**a**) and outside of the Europe (**b**). Dormant twig samples were collected from 155 tree species at 51 locations in 32 countries. Multiple locations within a country are merged and information is presented per country for better visibility. Different colours indicate fractions of samples belonging to different host genera. Maps were created using R 4.0.3 (R Core Team 2020. R: A language and environment for statistical computing. R Foundation for Statistical Computing, Vienna, Austria. https://www.R-project.org/).

## Results and discussion

The effects of local climate-, host- and geography-related factors on incidence-based β-diversity were similar within and across all tree-associated taxa (Fig. 2 a), indicating they jointly determine the presence of herbivorous insects and fungi associated with trees. Recent studies show the importance of host-phylogeny in shaping communities of tree-associated taxa^19,36^, but they underestimate the importance of climatic and geographic factors in structuring those communities. Our results thus highlight the value of high-resolution data for adequately estimating the relative importance of local environmental conditions and distance between sampling locations vs. host-related variables. Similar effects were found for the species turnover component of β-diversity (Simpson’s dissimilarity measure; Supplementary Fig. S1 a) indicating that turnover in species composition rather than species richness drove the responses. Overall, our models generally explained around 20% of the deviance in non-abundance weighted turnover (Fig. 2 a and Supplementary Table S1), showing that our measures of climate, host and geography are major drivers of tree associated communities. The general patterns remained largely consistent when samples containing no herbivorous insects or fungi were included (Supplementary Fig. S2 a and Supplementary Table S2), suggesting that the same drivers were important for infestation as for changes in community composition. While the size of the effects remained roughly equal for the abundance-weighted as for incidence-based herbivorous insect β-diversity (Fig. 2, Supplementary Fig. S1 and Supplementary Table S1), host-related factors had larger effects on fungal abundance-weighted β-diversity, in particular of saprotrophic fungi (Fig. 2 and Supplementary Fig. S1). Herbivorous insects therefore only seem able to occur in association with suitable hosts^42^, while fungi can be present in a wide range of hosts but can only become abundant in the most suitable hosts^22^.

**Figure 2.**
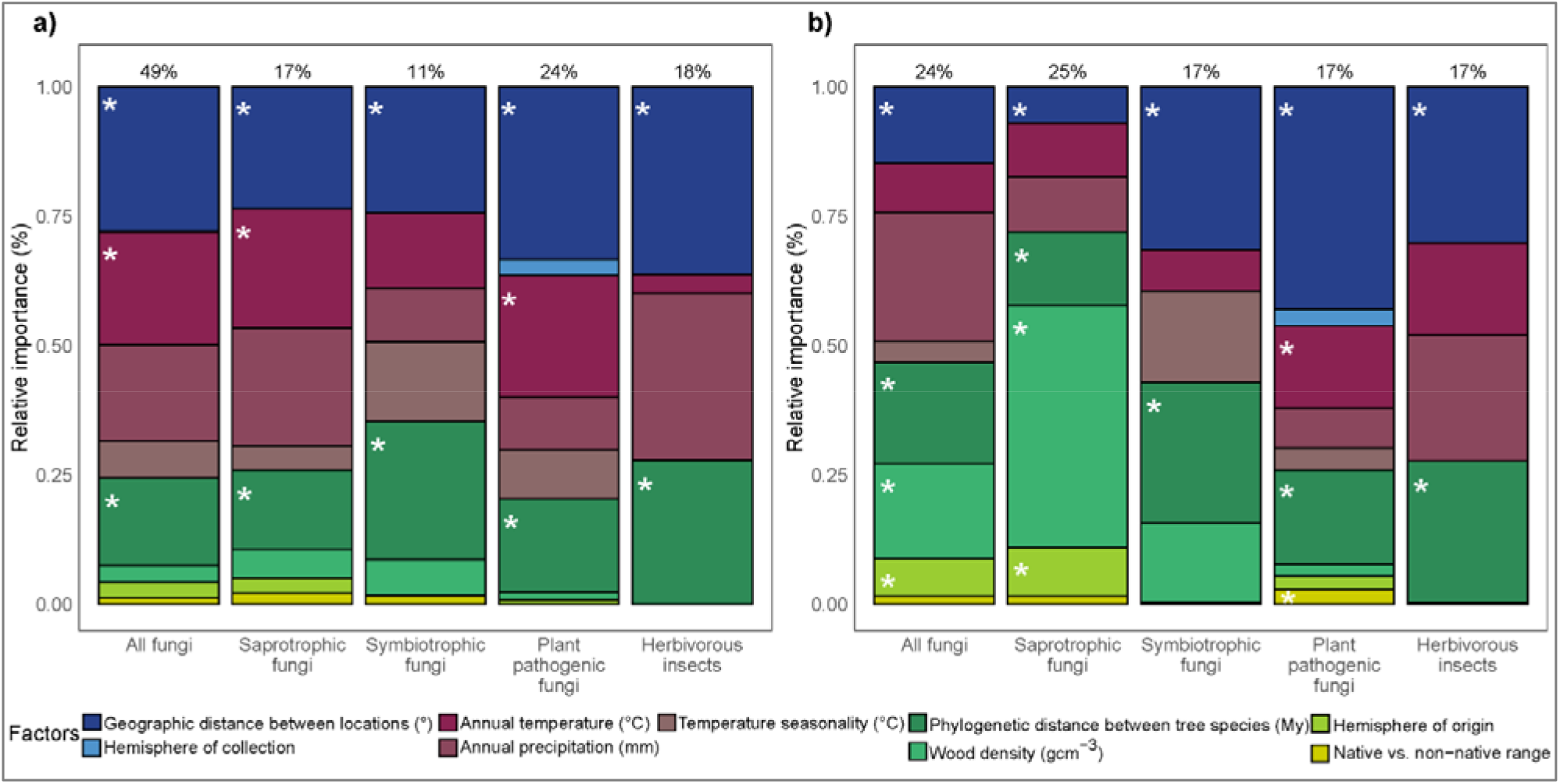
The relative importance of different variables for β-diversity of tree-associated fungi and herbivorous insects. The effects of variables on incidence-based (**a**, Sørensen) and abundance-weighted (**b**, Morisita Horn) β-diversity as assessed with generalised dissimilarity models. Geographic, climatic and host-related variables are shown in different shades of blue, red and green, respectively. The results are shown for all fungi together (N = 352), and for saprotrophic (N = 352), symbiotrophic (N = 223) and plant pathogenic fungi (N = 347) separately, and for herbivorous insects (N = 96). Numbers above bars indicate percent of total deviance explained by a model. The relative importance of variables in explaining the dissimilarities is calculated from max values of curves generated from generalised dissimilarity models. Significant factors (p < 0.05) are indicated with asterisks. Variable significance testing is done using 50 permutations. The results are shown for the entire data set and samples that contained insect and fungi (**“main analysis”**).

### Climatic factors

Climatic factors were one of the major drivers of β-diversity for fungi (especially saprotrophic and plant pathogenic fungi) and herbivorous insects (Fig. 2 and Supplementary Fig. S1). This supports previous studies, which showed that climatic factors, such as mean annual temperature, mean annual precipitation and temperature seasonality are strong drivers of the community composition of foliar fungal endophytes^17^ and insects^18^. High temperatures, especially above 10°C, had an increasing effect on incidence-based β-diversity (and turnover) of all, saprotrophic and plant pathogenic fungi (Fig. 3 a and Supplementary Fig. S3 a; GDMs test the size of the effect that variable has on β-diversity – increasing curve suggests an increasing effect of a variable on β-diversity) and abundance-weighted β-diversity of fungal plant pathogens (Fig. 3 b and Supplementary Fig. S3 b), but had no effect on herbivorous insect β-diversity. Similarly, mean annual precipitation was an important driver of turnover in all, saprotrophic, plant pathogenic fungi and herbivorous insects, especially above 1200 mm annual precipitation (Supplementary Fig. S3 a). Large differences in fungal and herbivorous insect communities among sites with high temperatures and precipitation suggest that growing in more extreme environmental conditions requires specialized traits and that tolerance of extreme climates trades off with competitive ability^43^, leading to restricted ranges for tolerant organisms. Tree-associated communities in more extreme environments, especially fungi in warmer and wetter areas, and herbivorous insects in wetter areas, might therefore be particularly susceptible to climate change since our results indicate that even small increases in temperature or precipitation may radically alter their composition. In addition, our results suggest that direct effects of climate change and shifts in tree species composition will have similarly large effects on herbivorous insect and fungal community composition, but that changes in tree species composition are likely to be particularly important in affecting dominant fungi.

**Figure 3.**
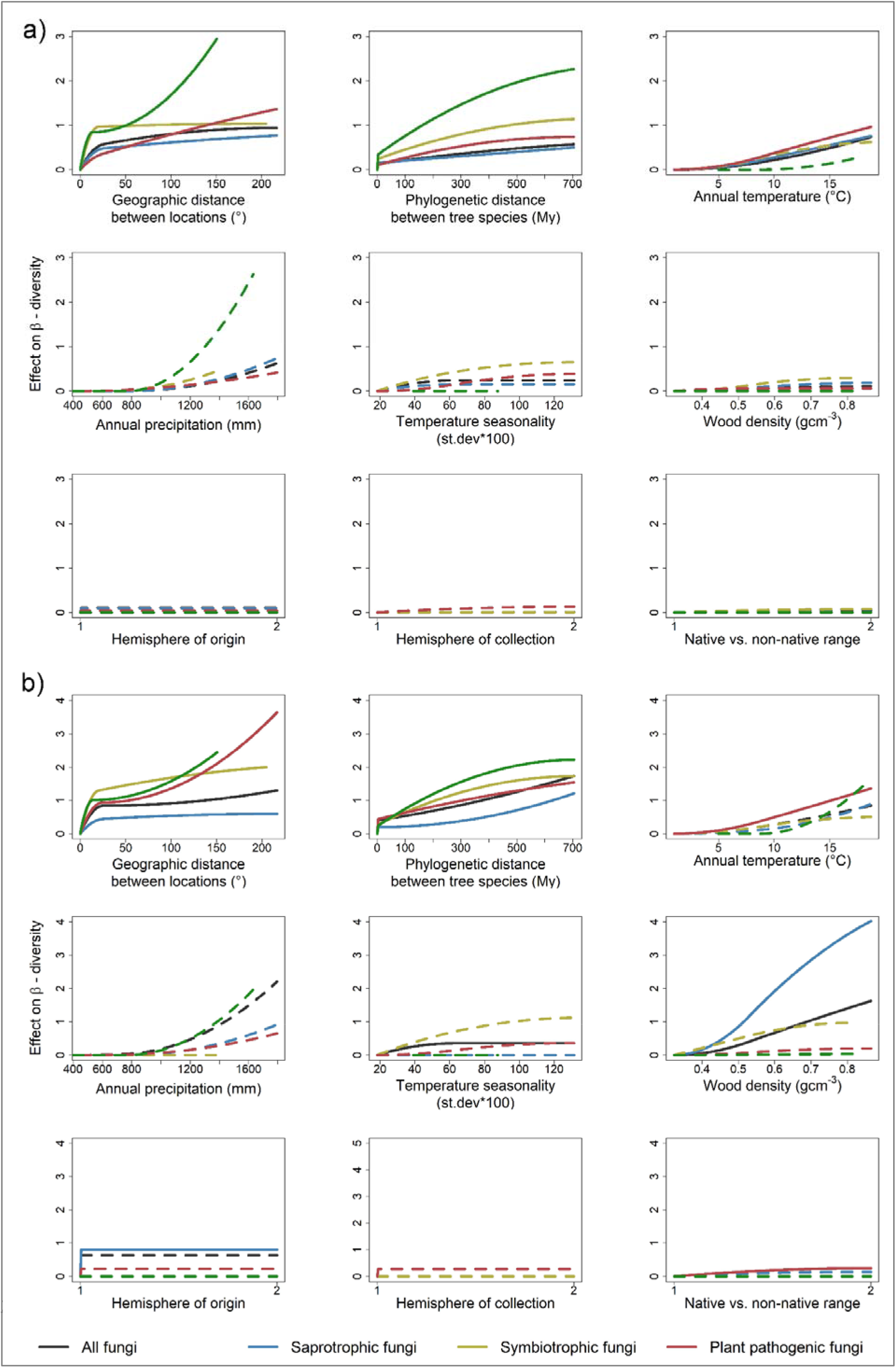
Effects of different variables on β-diversity of tree-associated fungi and herbivorous insects. The shape of the curve indicates the change in the effect of a variable on incidence-based (**a**, Sørensen) and abundance-weighted (**b**, Morisita Horn) β-diversity, at different points along the gradient of the variable. Generalized dissimilarity models were used to estimate these non-linear effects of the variables on β-diversity. The results are shown for all fungi together (N = 352), and for saprotrophic (N = 352), symbiotrophic (N = 223) and plant pathogenic fungi (N = 347) separately, and for herbivorous insects (N = 96). The final height of the curve indicates the relative importance of a variable in driving β-diversity. Significant factors (p < 0.05) are indicated with solid lines. Variable significance testing is done on a basis of 50 permutations. The results are shown for the entire data set and samples that contained insect and fungi (**“main analysis”**).

### Host related factors

Herbivorous insect and fungal communities were differentiated along a gradient of phylogenetic distance between tree species, with closely related hosts sharing more species than distantly related hosts (Fig. 3). Such patterns have been found for both fungi and insects^44^ and are expected from co-evolution. We observed initial steep increases in incidence-based and abundance-weighted β-diversity of all groups at very small phylogenetic distances, which corresponds to differences between conspecific (0 distance) and heterospecific tree hosts (>0 distance), and then a more or less linear increase (Fig. 3). The linear increase in compositional turnover with phylogenetic distance between hosts is similar to the relationship found in a study of phyllophagous beetles from tropical forest canopies^45^, but differs from studies showing exponential declines in the probability of spillovers of plant pathogenic fungi and insects between hosts with increasing phylogenetic distance^44^. Symptomatic plant pathogenic fungi and feeding herbivorous insects may therefore show greater host specialisation than non-symptomatic fungi and resting insects. Nevertheless, our results highlight the importance of phylogeny in predicting how likely tree species are to share fungi and insects and indicate the increased risk of host jumps and establishment of non-native organisms if closely related tree species are present in the new environment.

Fungal abundance-weighted β-diversity responded strongly to wood density, especially for saprotrophic fungi (Fig. 3 b and Supplementary Fig. S3 b). The rate of change accelerated above 0.5 g cm^-3^, which roughly corresponds to the difference between gymnosperm and angiosperm wood densities. Thus, our study confirms that the dominant fungi differ between and within angiosperms and gymnosperms^46,47^ and suggests that this is largely driven by the differences in wood density of the host. However, the significant effect of phylogeny in addition, indicates that host traits other than wood density also affect community composition of saprotrophic fungi, but complete data sets on host traits other than wood density are still unavailable and thus impossible to be studied.

Hemisphere of origin of host species was an important driver of abundance-weighted β-diversity of all and saprotrophic fungi (Fig. 2 b and Supplementary Fig. S1 b), but not of herbivorous insects, indicating that southern and northern hemisphere trees have distinct fungal communities. Separate co-evolutionary histories of plant lineages and associated organisms between two hemispheres^29,30^, due to unique geographic structures and climatic histories, might have resulted in the formation of specific co-adaptive traits of these organisms. Thus, the differences in fungal communities between tree species native to different hemispheres (but growing in the same hemisphere), confirm the tight relationships between tree species and associated organisms.

β-diversity of fungi generally did not differ between native or non-native tree species (no effect of native vs. non-native range; Fig. 2), the one exception being the abundance-weighted β-diversity of plant pathogenic fungi (Fig. 2 b). Similarly, no differences in β-diversity of herbivorous insects were found between native and non-native hosts, independent of the weighting of rare and abundant species, but our analyses suggest lower infestation incidence in non-native than in native trees (Supplementary Fig. S2), probably because non-native trees were released from their native enemies and did not recruit many new pests in the non-native range^32^. Native and non-native tree species sampled at the same site had similar insect and fungal species, probably because trees recruit organisms from closely related hosts present in the botanical gardens and arboreta that we sampled, although this could also be linked to sampling being done in botanical gardens or arboreta. In botanical gardens and arboreta many native and non-native tree species are grown under more or less standard conditions. This breaks up correlations between climate or soil and tree species composition, allowing us to better assess independent effects of host traits and environment on tree associated taxa. However, in botanical gardens and arboreta only a few trees of a species are grown in open areas which may result in high fragmentation of the habitat and altered micro-climatic structure (i.e., warmer and drier climate) in these locations in comparison with natural forests, as it is often the case for urban vs. non-urban sites^48^. This could reduce the effect of environmental filtering and increase the importance of colonization of generalist species from the surrounding matrix on insect and fungal assemblages in urban vs. non-urban sites^49^. Consequently, native trees in arboreta and botanical gardens may harbour a limited number of insects and fungi compared to trees in natural forests, potentially reducing differences with non-native taxa. The importance of the surrounding environment is highlighted in another study - in environments with native *Pinus sylvestris* stands, non-native *Pinus contorta* readily associates with fungal taxa already present, while in environments with distantly related *Nothofagus* species, *P. contorta* becomes associated with a unique fungal community that is more similar to fungal communities from the native range of western north America^50^. In addition, our results show that the fungal communities in native and non-native hosts were dominated by different species, possibly because non-native plants accumulate generalist but not specialist plant pathogenic fungi over time^33^. Taken together our results show that associated taxa are typically not specific to particular tree species and are rather shared between related hosts with similar traits and evolutionary history.

### Geographic factors

Geographic distance between locations was an important driver of incidence-based and abundance-weighted β-diversity of all groups of tree-associated organisms (Fig. 2). Climatic factors were included in our models, and hemisphere of collection was not an important driver of β-diversity in our study (Fig. 2 and Supplementary Fig. S1), suggesting that the effects of geographic distance between locations may be driven by dispersal limitation. Unmeasured environmental factors could also be important but at these large scales geographic barriers that limit the spread of organisms^31^, and inherent organism traits that affect the dispersal^15^ may have played a larger role. These results therefore show that microbial groups such as tree associated fungi do show geographic structure and probably dispersal limitation.

The shape of the geographic distance curve for all fungi (and saprotrophic and symbiotrophic fungi in particular) indicates a steep increase in incidence-based and abundance-weighted β-diversity only at intracontinental distances, i.e., up to around 15° (1,500 km; Fig. 3 and Supplementary Fig. S3). Very high β-diversity values were already reached at these distances, meaning that β-diversity could not increase further between continents. Previous studies report contradictory findings with some finding strong effects of geographic distance between locations^34,51,52^ on dissimilarities among insect and fungal communities and others finding no effect^19,36,53^. However, none of these studies included a continuous gradient from small to intercontinental distances, and our more extensive sampling provides strong evidence for relatively small scale geographic structure in tree associated fungi and herbivorous insects. The geographic separation of herbivorous insect and fungal communities within continents suggests that even the exchange of plant material across small spatial scales could lead to movement of associated herbivorous insects and fungi to areas where they previously did not occur.

Although geography was similarly important overall, plant pathogenic fungi and herbivorous insects showed a different pattern of geographic structure. The slope of the incidence-based β-diversity curve in communities of plant pathogenic fungi was relatively constant across the entire range and did not feature an initial steep increase (Fig. 3 a), suggesting better dispersal of plant pathogenic than other fungi. Many plant pathogenic fungi might be adapted to long distance aerial dispersal, to ensure regular reestablishment of diseases^54^, unlike wood decay fungi or mycorrhizal fungi, several of which show strong dispersal limitation even at small scales^55,56^. The effect of geographic distance on plant pathogenic fungi abundance-weighted β-diversity increased again when geographic distances between locations exceeded 100° (10,000 km; Fig. 3 b), which roughly corresponds to the average distance between continents. The same pattern was found for incidence-based β diversity of herbivorous insects (Fig. 3 and Supplementary Fig. S3). This indicates that communities of plant pathogenic fungi and insect herbivores on different continents are dominated by different species. Hence, further intercontinental exchange of plant material might facilitate new introductions of both insect and fungal pests.

### Conclusion

Our analyses of fungi and herbivorous insects, collected from trees growing on six continents, revealed the structure of potential pest-host associations across large spatial and phylogenetic scales. The results demonstrate the important roles of climate, host-related factors, and geographic location in jointly structuring herbivorous insect and fungal communities associated with trees. Although differences in climatic conditions strongly affected herbivorous insect and fungal communities, the importance of particular climatic drivers varied between functional groups, highlighting the importance of comparing across multiple tree-associated organisms. Climate change will, therefore, both directly and indirectly (through host range shifts) affect herbivorous insect and fungal communities. The specialisation of herbivorous insect and fungal communities on closely related and functionally similar trees indicates that new herbivorous insects and fungi are likely to be introduced as their host trees are moved, which calls for measures to reduce the likelihood of introducing the most harmful non-native organisms. This is particularly true for abundant fungal taxa. Safeguarding tree-derived environmental and societal benefits will, therefore, depend on limiting the establishment of new forest pests and increasing the resilience of trees and forest ecosystems to climate change.

## Methods

The sampling approach, identification of fungi and herbivorous insects and the molecular methods have been described in detail in Franić et al. (2022)^40^. Below, we provide a summarized version and full details of the statistical analyses presented in this manuscript.

### Field collection

Dormant twigs were collected from 155 tree species at 51 locations in 32 countries, in the northern and southern hemispheres. Very few tree genera occur naturally in both hemispheres (e.g., *Podocarpus*), so we selected 17 genera that occur widely across either the northern or southern hemisphere. We sampled in botanical gardens and arboreta and we purposefully sampled native and non-native congeneric or confamiliar tree species at each location. Formal tree species identification was conducted by the co-authors and expert personnel in botanical gardens and arboreta where samples were collected^38^. Voucher specimens of the material were not deposited in a publicly available herbarium because our methods for herbivorous insect and fungal assessment were destructive.

One sample consisted of twenty 50 cm long asymptomatic twigs, collected from up to 5 individual trees per species, at each location. Twigs were collected in the month with the shortest day-length in the year (end of December 2017 in the Northern hemisphere, end of June 2018 in the Southern hemisphere). Eleven samples, from a tropical region in Tanzania, were collected in June 2018. Dormant tree twigs were sampled in winter in order to accurately assess herbivorous insect and fungal species which are likely to be moved with traded plants as most woody plants are traded in winter to be planted the following spring, and to reduce the risk of introducing foliar pests with deciduous trees. Evergreen tree species, which were also considered, were sampled with leaves/needles because they do not lose foliage during winter, and are thus sold with leaves/needles. Although plants are sometimes traded with roots for species which can’t be grown from twigs (i.e., budwood), the roots were not sampled in this study because this would have been highly destructive to the plants.

Fungi were assessed from a total of 352 samples from 145 native and non-native tree species, belonging to nine families of angiosperms and gymnosperms from 44 locations in 28 countries on five continents. Herbivorous insects were assessed from 227 samples of 109 tree species, collected at 31 locations and in 18 countries in two hemispheres. At some locations it was not possible to assess both herbivorous insects and fungi because of the lack of expertise and/or infrastructure to collect and/or identify specimens belonging to both groups.

### Fungal assessment

After surface sterilization and air drying on a sterile bench, the following material from each of 20 twigs per sample was pooled: half of one bud, a 0.5 cm long piece of a needle (from gymnosperms) or a 0.25 cm^2^ leaf (for evergreen angiosperms) and a 0.5 cm long piece of twig.

#### DNA extraction, PCR amplification and Illumina sequencing

DNA was extracted from 50 mg of pooled and ground tissue. DNA concentrations were quantified and DNA was diluted to 5 ng/μl. The internal transcribed spacer (ITS2 region) of the ribosomal operon was amplified as described in Franić et al. (2019). Each sample was amplified in triplicates which were then pooled. Successful PCR amplification was confirmed by visualization of the PCR products. Pooled amplicons were sent to the Génome Québec Innovation Centre at McGill University (Montréal, Quebec, Canada) for barcoding using the Fluidigm Access Array technology (Fluidigm, South San Francisco, CA, USA) and paired-end sequencing on the Illumina MiSeq v3 platform (Illumina Inc., San Diego, CA, USA).

#### Bioinformatics and taxonomical classification of Amplicon sequence variants (ASVs)

Quality filtering and delineation into ASVs were done with a customized pipeline largely based on VSEARCH^57^, as described by Herzog et al. (2019). Taxonomic classification of ASVs was conducted using Sintax^59^ implemented in VSEARCH against the UNITE v.7.2 database^60^ with a bootstrap support of 80%.

#### Assignment of fungal trophic groups

Fungal ASVs were assigned to trophic groups using the FUNGuild tool ^61^. FunGuild consists of a community-annotated data base and a bioinformatic script that assigns fungal ASVs to trophic groups (Pathotroph = “receiving nutrients at the expense of the host cells and causing disease”, Saprotroph = “receiving nutrients by breaking down dead host cells”, Symbiotroph = “receiving nutrients by exchanging resources with host cells”) based on taxonomic information. We only considered ASVs that were assigned to a single trophic group. Within pathotrophs we selected ASVs that were assigned to a “Plant Pathogen” guild. Throughout the manuscript we use terms: plant pathogenic, symbiotrophic and saprotrophic fungi. We only considered assignments that were ranked as probable and highly probable in FunGuild, as recommended by the authors^61^. Around 15% of total 12,721 fungal ASVs were assigned to a trophic group following our approach. Saprotrophic fungi were represented by 1,018 ASVs, plant pathogenic fungi by 754 ASVs and symbiotrophic fungi by 127 ASVs.

### Herbivorous insects

The collected twigs were brought to a laboratory and were first screened for insects that overwinter as adults. Twigs were then kept at room temperature with the cut ends immersed in water to allow the development of insects that overwinter as larvae, pupae or eggs. Twigs were inspected for insects daily for 4 weeks and all collected insects were stored in 95% ethanol for further examination.

#### Morphological and molecular identification

Insects were sorted in respect to their taxonomic orders and feeding guilds (i.e., herbivores, predators, parasitoids and other). Herbivorous insects were further sorted into morphospecies and at least one specimen per morphospecies was stored at -20°C for molecular analysis. Genomic DNA was extracted with a KingFisher (Thermo Fisher Scientific) extraction protocol suitable for insects in a 96-well plate. PCR for the COI was carried out in 25 µl reaction volume as described in Franić et al 2019. The success of amplification was verified by electrophoresis of the PCR products. A standard Sanger sequencing of the PCR products in both directions with the same primers was done at Macrogen Europe, Amsterdam, Netherlands. Sequences were assembled and edited with CLC Workbench (Version 7.6.2, Quiagen) and compared to reference sequences in BOLD^62^ or the National Centre for Biotechnology Information (NCBI) GenBank databases and matched to the best matching reference sequence for identification. Pictures of insects with unclear identification results were sent to experts for further examination.

Since the identification of Thysanoptera was not clear for all specimens, a phylogenetic analysis was conducted with MEGA6^63^ software using the General Time Reversible substitution model as described in Franić et al. (2022)^40^.

### Sample metadata

Climate data were obtained from the WorldClim database^64^ at a resolution of 2.5 min and represent the average for the years 1970-2000.

A host-tree phylogeny was constructed with the phylomatic^65^ function from the package brranching^66^ in R^67^ using the “zanne2014” reference tree^68^. One *Eucalyptus* sample collected in Argentina and two *Eucalyptus* samples collected in Tunisia were not identified to species. Therefore, to place them in the phylogeny, we assigned them to congeneric species that were not sampled in this study but that we considered as representative samples of phylogenetic diversity from across the *Eucalyptus* genus (*Eucalyptus viminalis, Eucalyptus robusta* and *Eucalyptus radiata*). Pairwise phylogenetic distances between study tree species were calculated using the cophenetic function^67^.

Wood densities for study species were retrieved from the Global Wood Density Database^69^. If more than one measure was available for a given species in the data base, the mean value was used. If there were no wood density data available for a tree species (61 out of 155 tree species), we used the mean wood density for its genus (60 species), or if no congeneric species were present, for its family (1 species).

### Statistical analyses

We analysed three measures of β-diversity based on Hill numbers^70^, in which increasing weight is given to species abundances. The three β-diversity measures were the Sørensen index (q = 0), Horn’s index (q=1), and Morisita–Horn index (q = 2)^41^. Since these are all similarity indices, we converted them to dissimilarity measures as 1-similarity index. In the text we refer to Sorensen index (q = 0) as incidence-based β-diversity and Horn’s (q=1) and Morisita-Horn index (q = 2) are referred to as abundance-weighted β-diversity. Additionally, to test for effects on β-diversity that are due to pure turnover and not differences in species richness differences, we used the Simpson dissimilarity (species turnover), which is the turnover component of Sørensen index. This was calculated with the function beta.pair from the betapart package^71^.To analyze the non-linear response of fungal and herbivorous insect β-diversity to differences in geographic, host and climate related factors, we used generalized dissimilarity models (GDM)^72^. GDMs allowed us to assess the relative importance of different factors in explaining turnover in herbivorous insect and fungal community composition, while keeping all other factors constant. They also allowed us to test how the effects of an environmental variable on turnover changed along the environmental gradient, e.g., to test whether small or large differences in a climatic variable caused the most turnover in tree associated communities. We included spatial distance between sites (i.e., “geographic distance between locations”) and differences in hemisphere of collection (binary variable: same hemisphere or different hemispheres) as measures of geographic differences, and differences in mean annual temperature, mean annual precipitation and temperature seasonality between sites as measures of climatic differences. Phylogenetic distance between tree species, differences in wood density, differences in hemisphere of origin (i.e., the hemisphere to which the tree species are native) and differences in native vs. non-native range, i.e., native or non-native status of the tree species at the sampling location (the latter two as binary variables) were included to estimate the effect of differences in host factors on β-diversity. Although GDMs assume that environmental predictors are continuous variables or, at least, that these variables consist of ordered categories, categorical variables can also be used in GDMs^72^. For incorporating our categorical variables (i.e., hemisphere of origin, hemisphere of collection and native vs. non-native range) into GDMs we assigned each pair of sites a distance of one if they occur in the same class, or two if they occur in different classes, and then treat this binary distance measure in the same manner as other distance data. We fitted GDMs for herbivorous insects, all fungi together and saprotrophic, symbiotrophic and plant pathogenic fungi separately. For herbivorous insects, the hemisphere of collection was not included because we only sampled insects from the northern hemisphere. GDMs were fitted using the gdm function from the *gdm* package^73^ in R. Matrix permutations were used to perform model and variable significance testing and to estimate variable importance using the function gdm.varImp^22^ with default parameters and 50 permutations.

To account for differences in sequencing depth across samples, which is purely a methodological artefact, sequence data were normalized to the same number of reads per sample. When data were rarefied to 10,000 reads per sample, the general patterns remained similar (Supplementary Fig. S4 and S5 and Supplementary Table S3).

The main analysis considered samples that contained herbivorous insects or fungi of different guilds because similarity indices cannot be calculated for empty samples. We detected fungi in 352 samples and all of those contained saprotrophic fungi, however symbiotrophic and plant pathogenic fungi were absent from 129 and 5 samples, respectively. Herbivorous insects were not detected in 127 out of 217 samples. This means that different models for the different groups were fitted to different numbers of plots. To include the blank samples in the analysis, which allowed us to assess the drivers of infestation, we calculated Sørensen dissimilarity by adding dummy species to the datasets (“zero-adjusted analysis”; Supplementary Fig. S2 a and Supplementary Table S2). This approach was previously shown to improve interpretability of results^74^. However, it was not used for abundance weighted β-diversity measures as it is only appropriate for dissimilarity measures from the “Bray–Curtis family”. The general patterns of β-diversity remained largely consistent when samples containing no herbivorous insects or fungi were included (Supplementary Fig. S2 a), suggesting that the same drivers were important for infestation as for changes in community composition. However, geographic factors were not important drivers of differences in infestation by symbiotrophic fungi and herbivorous insects (Supplementary Fig. S2 a). This is because many samples contained no symbiotrophic fungi or herbivorous insects (i.e., 129 out of 325 and 121 out of 217, respectively) and these were evenly spread along the geographic gradient.

We were not able to assess both fungi and herbivorous insects simultaneously from all studied samples. To be able to directly compare the relative importance of different variables on β-diversity between herbivorous insects and fungi we therefore repeated the analysis considering samples from which both groups were assessed (167 samples). This approach restricted our data set to European and Siberian samples. We analyzed this data set by excluding blank samples (“overlap analysis”; for results see Supplementary Fig. S6-S7 and Supplementary Table S4) and using zero-adjusted Sørensen dissimilarity (“zero-adjusted overlap analysis”; for results see Supplementary Fig. S2 b and Supplementary Table S5). The results were broadly consistent with the results of the main analysis. However, the overall relative effects of host-related variables on β-diversity increased (Supplementary Fig. S6-S7), because the resolution and length of gradients of geographic distance between locations and climatic variables decreased (Supplementary Fig. S8). Mean annual temperature and mean annual precipitation were not significant drivers of β-diversity of tree-associated fungi and herbivorous insects in this analysis (Supplementary Fig. S6). This is because the smaller climatic variation in the overlap analysis, i.e., no high values above 12 °C and 1200 mm (Supplementary Fig. S8). Furthermore, the overlap analysis revealed a strong effect of hemisphere of origin on incidence-based and abundance-weighted β-diversity of herbivorous insects (Supplementary Fig. S6) suggesting specialization of herbivorous insects between hosts native to southern vs northern hemisphere, similar as it was shown for all and saprotrophic fungi in the main analysis.

## Data availability

The data used in this manuscript, as well as the detail methods on how they were collected and public repositories in which they are stored, are described in Franić et al. (2022)^40^. The raw paired-end Illumina sequencing reads of the ITS2 region are archived at the NCBI Sequence Read Archive under BioProject accession number PRJNA70814822. Assembled herbivorous insect COI sequences are deposited in GenBank database under accession numbers MW441337-MW441767.

## Code availability

R functions and databases used for generating the sample metadata are specified in the method section. A customized pipeline used for quality filtering of the raw sequence data obtained from HTS, delineation into ASVs and taxonomic classification of ASVs as described in Herzog et al. (2019) is available as a “ITS2.bash” file from the Zenodo repository^75^.

## Acknowledgments

We gratefully acknowledge the financial support of the Swiss National Science Foundation (Project C15.0081) and the Swiss Federal Office for the Environment (Grant 00.0418.PZ/ P193-1077). This work was supported by COST Action “Global Warning” (FP1401). CABI is an international intergovernmental organisation, and R.E., M.K., H.L. and I.F. gratefully acknowledge the core financial support from our member countries (and lead agencies) including the United Kingdom (Foreign, Commonwealth & Development Office), China (Chinese Ministry of Agriculture and Rural Affairs), Australia (Australian Centre for International Agricultural Research), Canada (Agriculture and Agri-Food Canada), Netherlands (Directorate General for International Cooperation), and Switzerland (Swiss Agency for Development and Cooperation). See https://www.cabi.org/aboutcabi/who-we-work-with/key-donors/ for full details. M.B. and M.K.H. were financially supported by the Slovak Research and Development Agency (Project APVV-19-0116). H.B. would like to thank the botanist Jorge Capelo who helped with Myrtaceae identification and INIAV IP for supporting her contribution to this study. Contributions of M. de G. and B.P. were financed through Slovenian Research Agency (P4-0107) and by the Slovenian Ministry of Agriculture, Forestry and Food (Public Forestry Service). G.C, C.B.E. and A.F.M. were supported by OTKA 128008 research grant provided by the National Research, Development and Innovation Office. Contributions of K.A. and R.D. were supported by the Estonian Science Foundation grant PSG136. M.J.J., C.L.M. and H.P.R. were financially supported by the 15. Juni Fonden (Grant 2017-N-123). P.B., B.G. and M.Ka. were financially supported by the Ministry of Science and Higher Education of the Republic of Poland for the University of Agriculture in Krakow (SUB/040013-D019). C.N. was financially supported by the Slovak Research and Development Agency (Grant APVV-15-0531). N.K. was partially supported by Russian Foundation for Basic Research (Grants 15-29-02645, 19-04-01029). R.OH. was supported by funding from DAERA, and assistance from David Craig, AFBI. T.P. thanks the South African Department of Forestry, Fisheries and the Environment (DFFE) for funding noting that this publication does not necessarily represent the views or opinions of DFFE or its employees. In preparing the publication, materials of the bioresource scientific collection of the CSBG SB RAS “Collections of living plants indoors and outdoors” USU_440534 (Novosibirsk, Russia) were used. M.Z. was financially supported by Ministry of education, science and technological development of the Republic of Serbia (Contract 451-03-68/2020-14/200197). We acknowledge the Genetic Diversity Centre (GDC) at ETH Zurich for providing computational infrastructure and acknowledge the contribution of McGill University and Génome Québec Innovation Center (Montréal, Quebec, Canada) for pair-end sequencing on Illumina MiSeq. The funders had no role in study design, data collection and analysis, decision to publish, or preparation of the manuscript.

## Author contributions

I.F., S.P. and R.E. designed the study. I.F. compiled the common sampling and laboratory protocols and coordinated sample collection. All co-authors did sample collection and/or DNA extractions for fungal assessment and/or insect rearing. I.F. finalized laboratory preparation of samples for HTS. M.H. supervised amplicon sequencing approach and performed the bioinformatic analyses of the HTS sequence data. N.F. and B.R. did morphological and molecular identification of insect with support of M.M.G.. I.F., E.A. and R.E. analysed the data and wrote the manuscript with contribution from all authors.

## Additional information

Supplementary information is available for this paper.

Correspondence and requests for materials should be addressed to I.F.

Plant material used in this study was obtained with permission and assistance of responsible personnel of botanical gardens and arboreta where samples were collected^38^.

All methods were carried out in accordance with relevant guidelines and regulations.

