## Supplemental Material for "Climate, host and geography shape insect and fungal communities of trees"

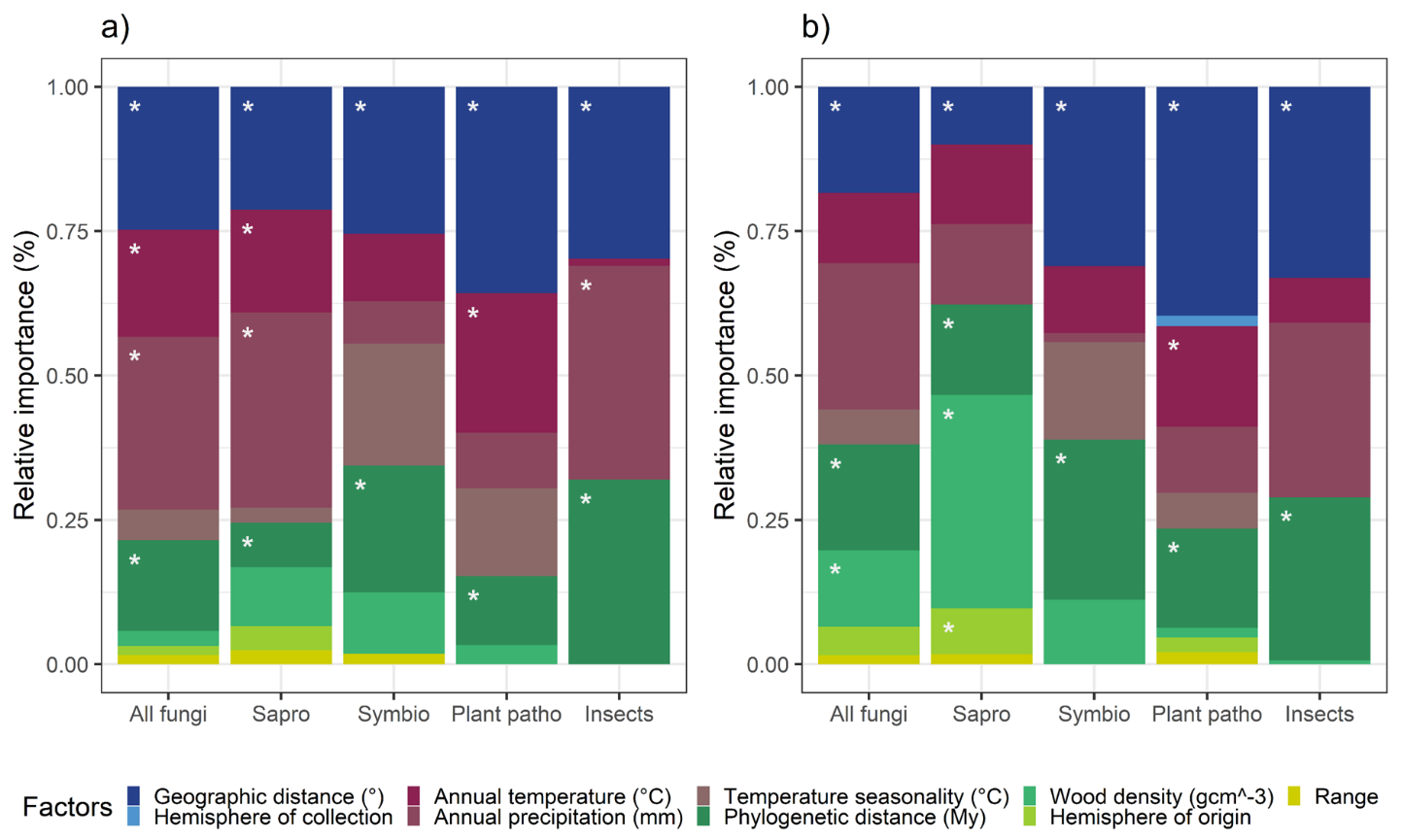

### **Supplementary Information**

Supplementary Figure S1. **The relative importance of different variables for β-diversity of tree-associated organisms.** **a**, **b** the effects of given variables on species turnover component of β-diversity (**a**, βsim) and abundance weighted β-diversity (**b**, Horn q=1) as assessed with GDMs. Geographic, climatic and host related variables are shown in different shades of blue, red and green, respectively. The results are shown for all fungi (“All fungi”), saprotrophs (“Sapro”), symbiotrophs (“Symbio”), plant pathogens (“Plant patho”) and herbivorous insects (“Insects”). Significant factors are indicated with asterisks. Variable significance testing is done on a basis of 50 permutations. The results are shown for the entire data set and samples that contained insect and fungi (**“main analysis”**).

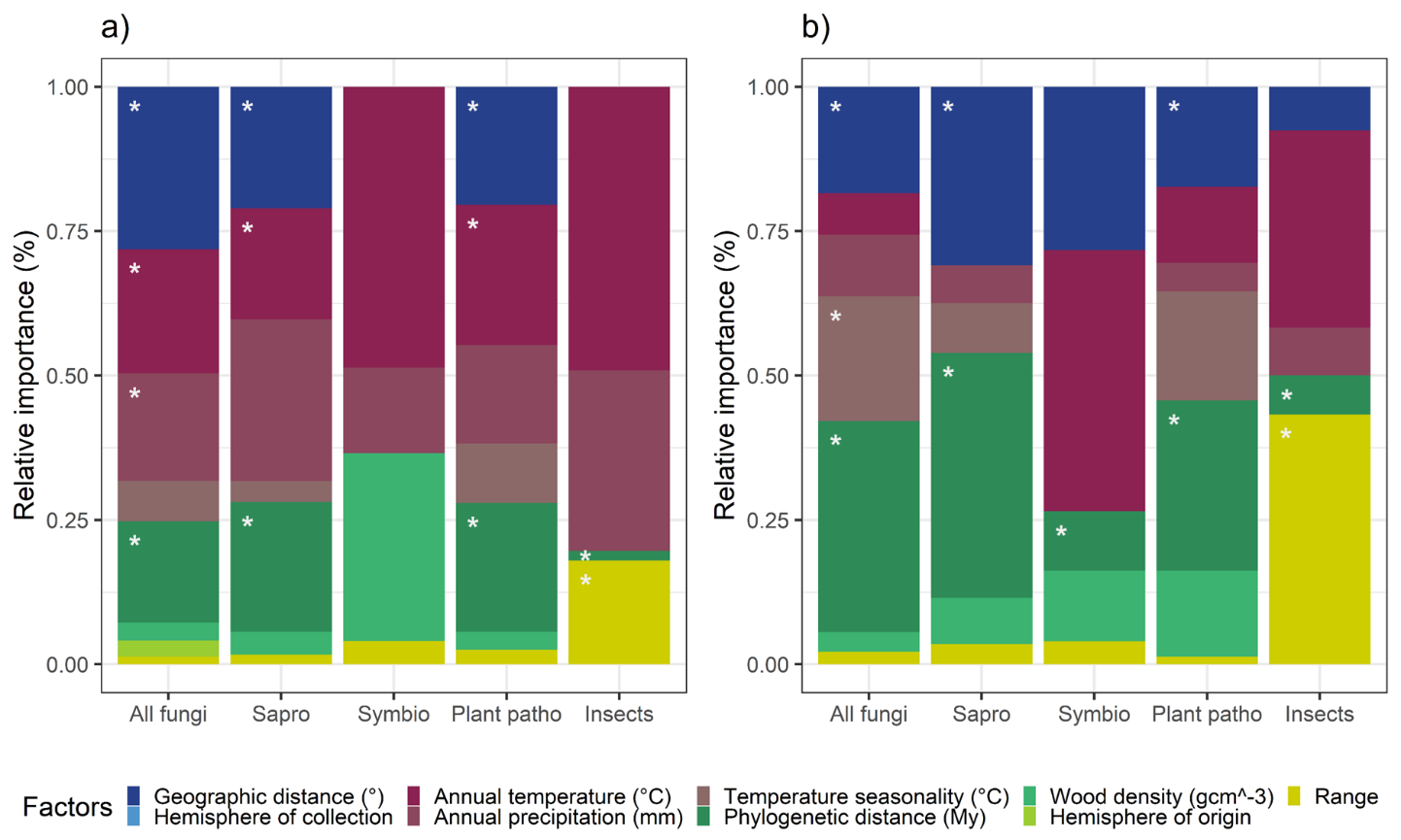
Supplementary Figure S2. **The relative importance of different variables for β-diversity of tree-associated organisms.** **a**, **b** the effects of given variables on incidence based β-diversity (Sørensen, q=0) as assessed with GDMs for zero adjusted data for the entire data set (**a**, “zero adjusted analysis”) and samples from which both insects and fungi were assessed (**b,** “overlap zero adjusted analysis”). Geographic, climatic and host related factors are shown in different shades of blue, red and green, respectively. The results are shown for all fungi (“All fungi”), saprotrophs (“Sapro”), symbiotrophs (“Symbio”), plant pathogens (“Plant patho”) and herbivorous insects (“Insects”). Significant factors are indicated with asterisks. Variable significance testing is done on a basis of 50 permutations.

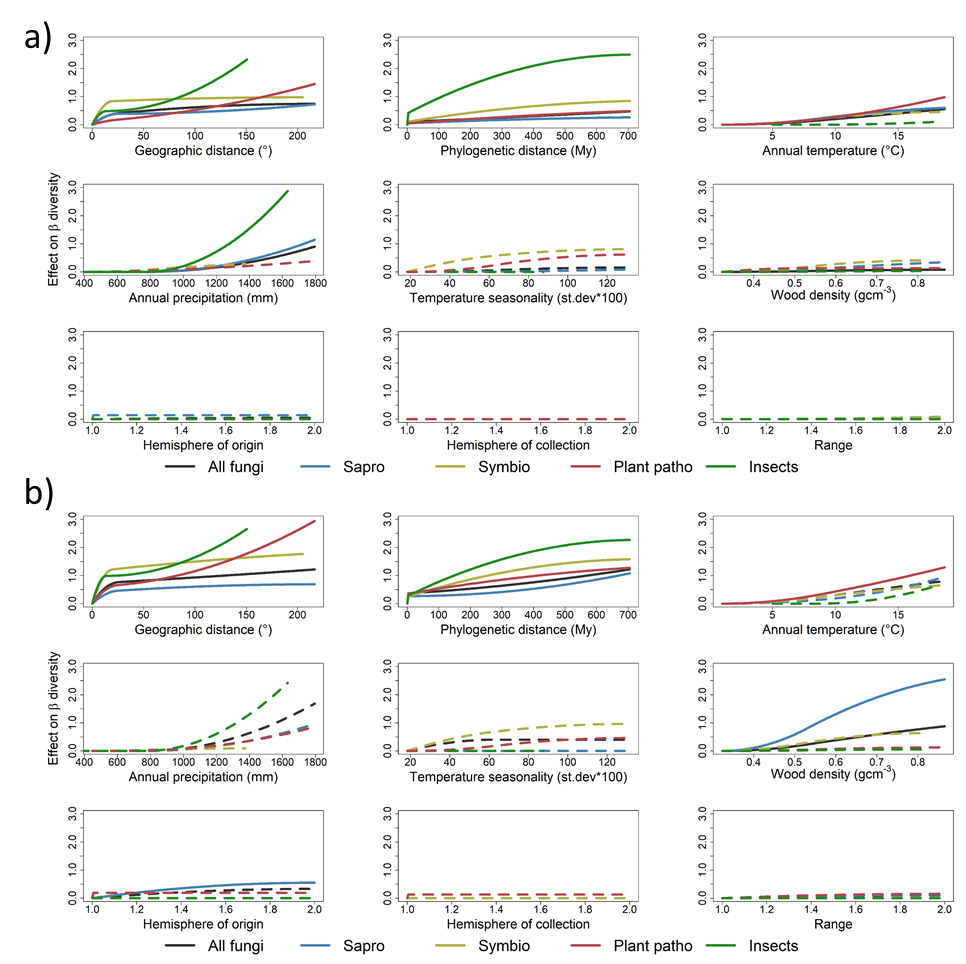
Supplementary Figure S3. **Effects of different variables on β-diversity of tree-associated organisms.** **a, b** The shape of the curve indicates the change in species turnover component of β-diversity (**a**, βsim) and abundance-weighted β-diversity (**b**, Horn q=1) along the variable gradient as assessed with GDMs. The results are shown for all fungi (“All fungi”), saprotrophs (“Sapro”), symbiotrophs (“Symbio”), plant pathogens (“Plant patho”) and herbivorous insects (“Insects”). The final height of the curve indicates the relative importance of a variable. Significant factors are indicated with solid lines. Variable significance testing is done on a basis of 50 permutations. The results are shown for the entire data set and samples that contained insect and fungi (**“main analysis”).**

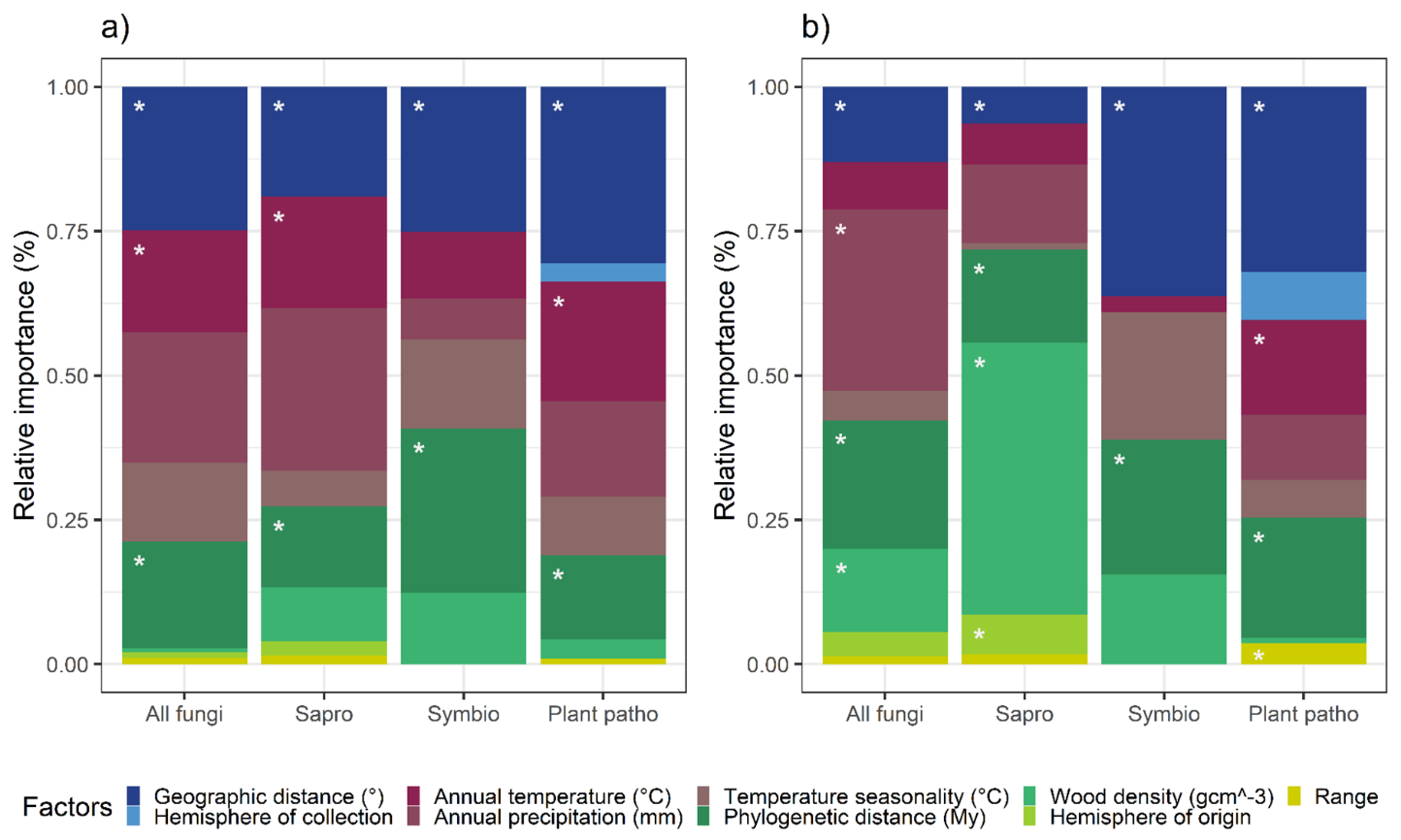
Supplementary Figure S4. **The relative importance of different variables for β-diversity of tree-associated fungi and insects.** The effects of variables on incidence-based (**a**, Sørensen) and abundance-weighted (**b**, Morisita Horn) β-diversity as assessed with generalised dissimilarity models. Geographic, climatic and host-related variables are shown in different shades of blue, red and green, respectively. The results are shown for all fungi together (“All fungi”), and saprotrophs (“Sapro”), symbiotrophs (“Symbio”) and plant pathogens (“Plant patho”) separately. Significant factors (p < 0.05) are indicated with asterisks. Variable significance testing is done using 50 permutations. The results are shown for the entire data set rarefied to 10,000 reads per sample and samples that contained insect and fungi (**“rarefied data analysis”**).

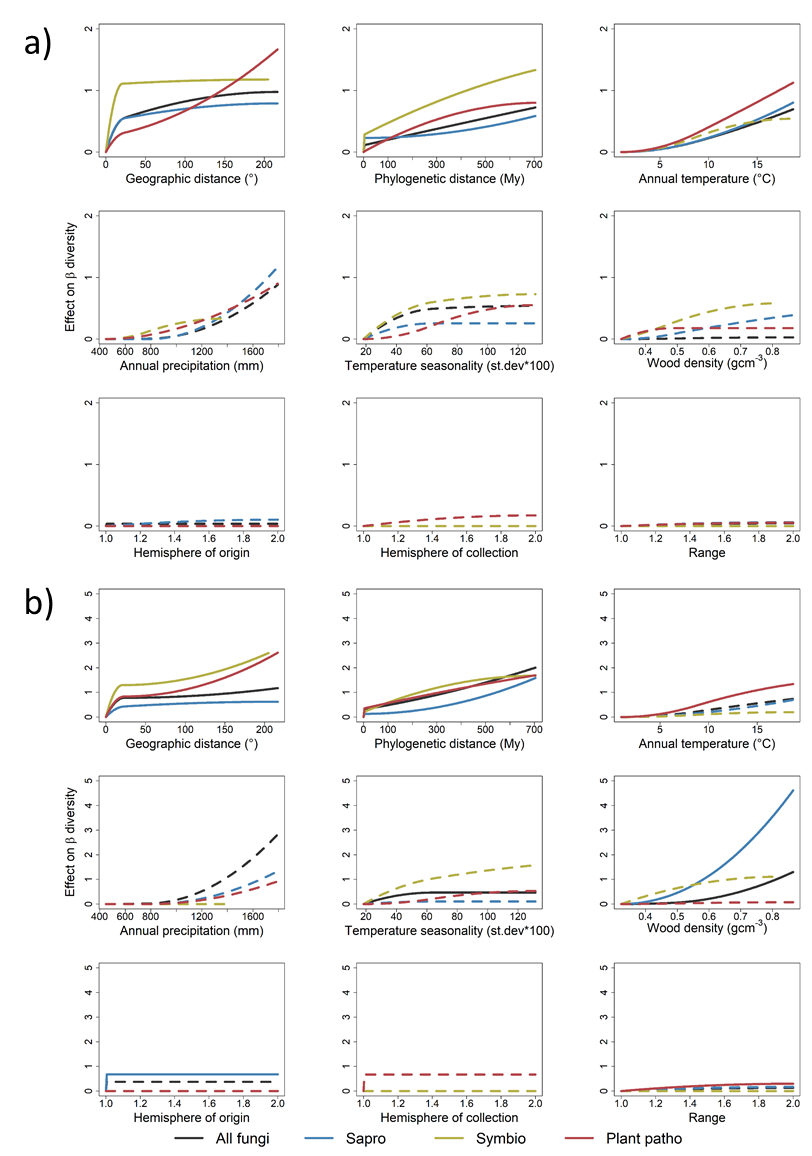

Supplementary Figure S5. **Effects of different variables on β-diversity of tree-associated fungi and insects.** The shape of the curve indicates the change in the effect of a variable on incidence-based (**a**, Sørensen) and abundance-weighted (**b**, Morisita Horn) β-diversity, at different points along the gradient of the variable. Generalized dissimilarity models were used to estimate these non-linear effects of the variables on β diversity. The results are shown for all fungi together (“All fungi”), and saprotrophs (“Sapro”), symbiotrophs (“Symbio”) and plant pathogens (“Plant patho”) separately. The final height of the curve indicates the relative importance of a variable in driving β-diversity. Significant factors (p < 0.05) are indicated with solid lines. Variable significance testing is done on a basis of 50 permutations. The results are shown for the entire data set rarefied to 10,000 reads per sample and samples that contained insect and fungi (**“rarefied data analysis”**).

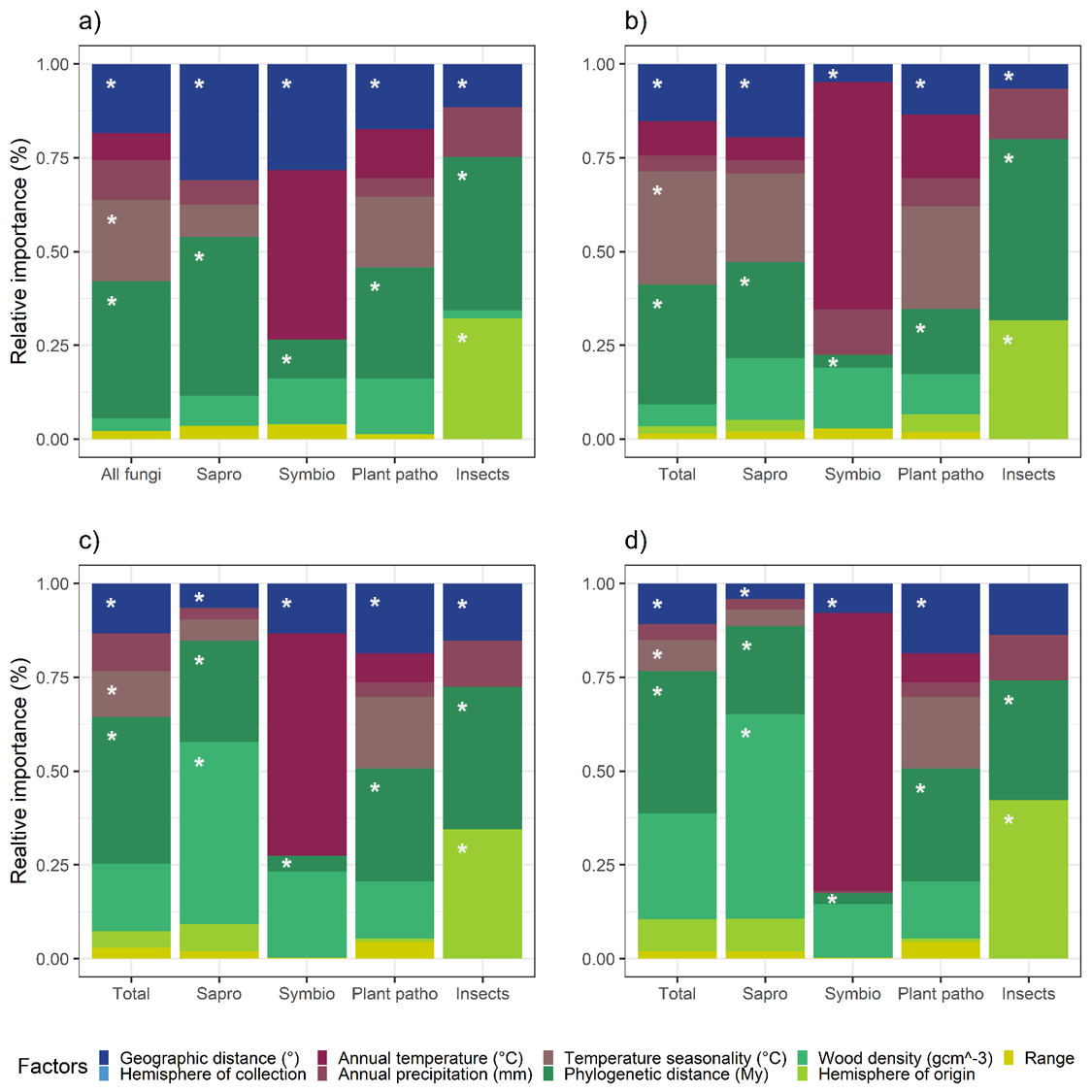

Supplementary Figure S6. **The relative importance of different variables for β-diversity of tree-associated organisms. a**, **b, c, d** the effects of given variables on incidence based β-diversity (**a**, Sørensen q=0), species turnover component of β-diversity (**b**, βsim) and abundance weighted β-diversity (**c,** Horn q=1; **d,** Morisita-Horn q=2) as assessed with GDMs. Geographic, climatic and host related factors are shown in different shades of blue, red and green, respectively. The results are shown for all fungi (“All fungi”), saprotrophs (“Sapro”), symbiotrophs (“Symbio”), plant pathogens (“Plant patho”) and herbivorous insects (“Insects”). Significant factors are indicated with asterisks. Variable significance testing is done on a basis of 50 permutations. The results are shown for the samples from which both insects and fungi were assessed and which contained insects and fungi (**“overlap** **analysis”**).

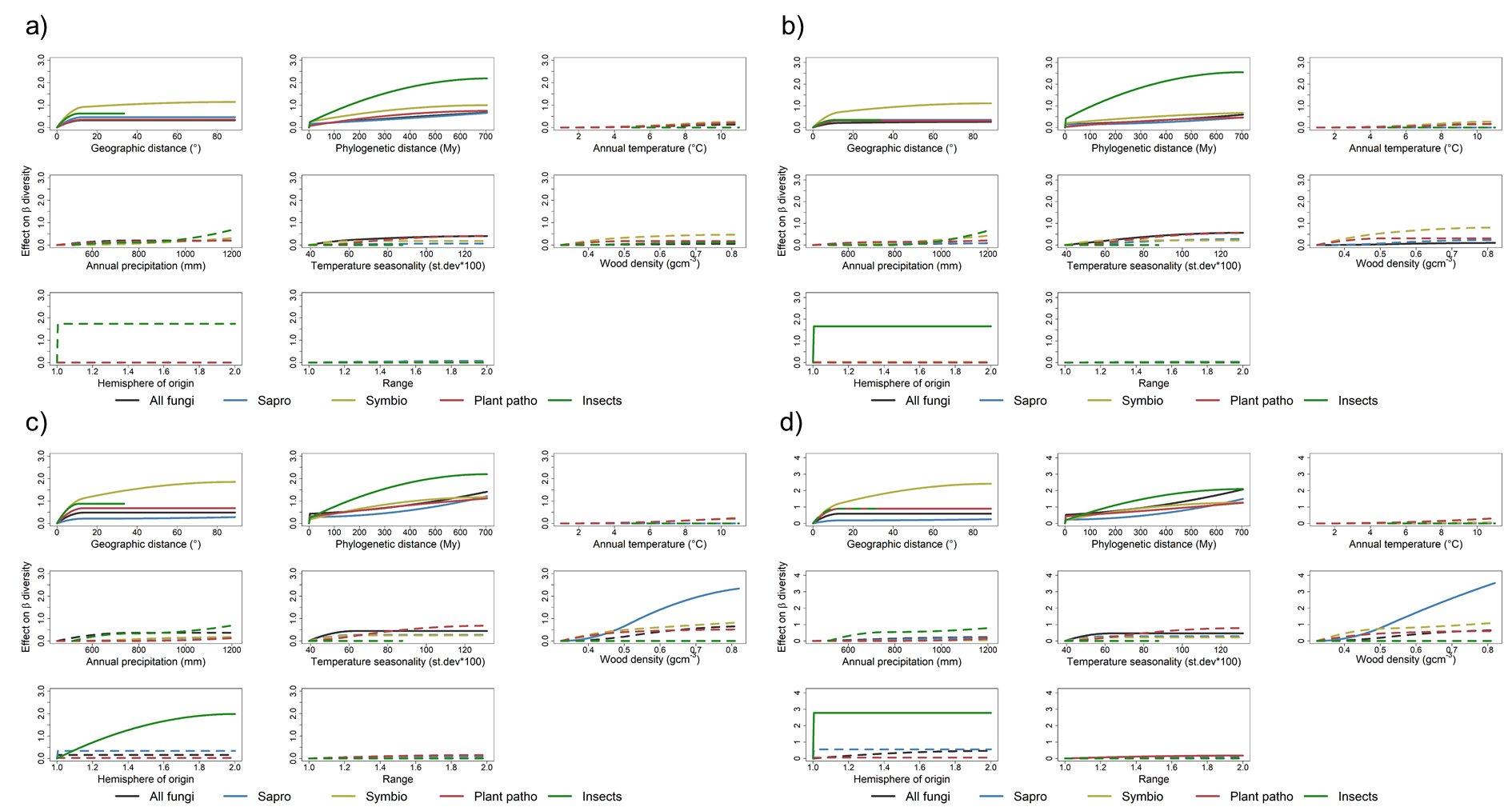

Supplementary Figure S7. **Effects of different variables on β-diversity of tree-associated organisms.** **a, b, c, d** The shape of the curve indicates the change in incidence based β-diversity (**a**, Sørensen q=0), species turnover component of β-diversity (**b**, βsim) and abundance weighted β-diversity (**c**, Morisita q=1; **d**, Morisita-Horn q=2) along the variable gradient as assessed with GDMs. The results are shown for all fungi (“All fungi”), saprotrophs (“Sapro”), symbiotrophs (“Symbio”), plant pathogens (“Plant patho”) and herbivorous insects (“Insects”). The final height of the curve indicates the relative importance of a variable. Significant factors are indicated with solid lines. Variable significance testing is done on a basis of 50 permutations. The results are shown for the samples from which both insects and fungi were assessed and which contained insects and fungi (**“overlap analysis”**).

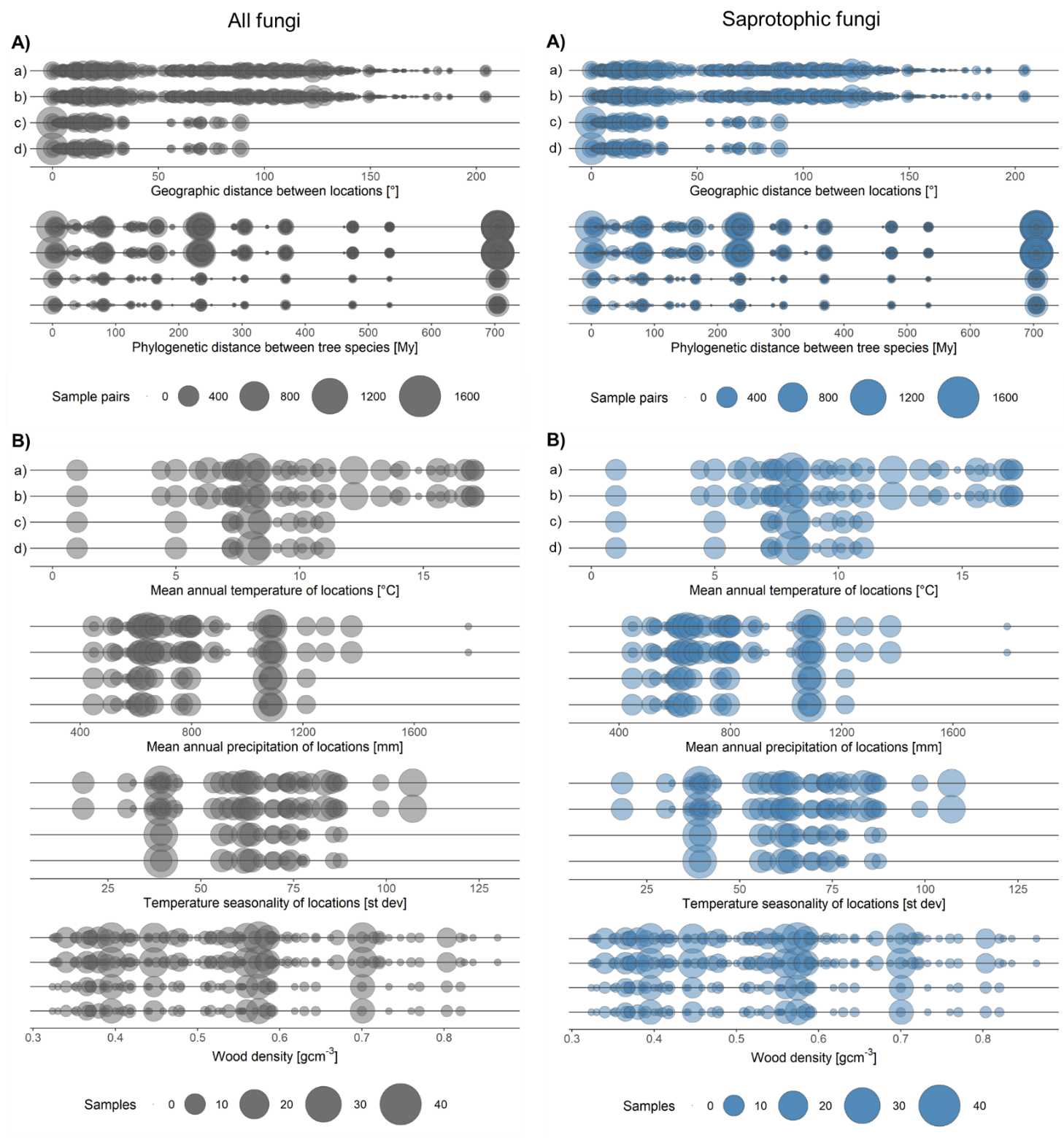

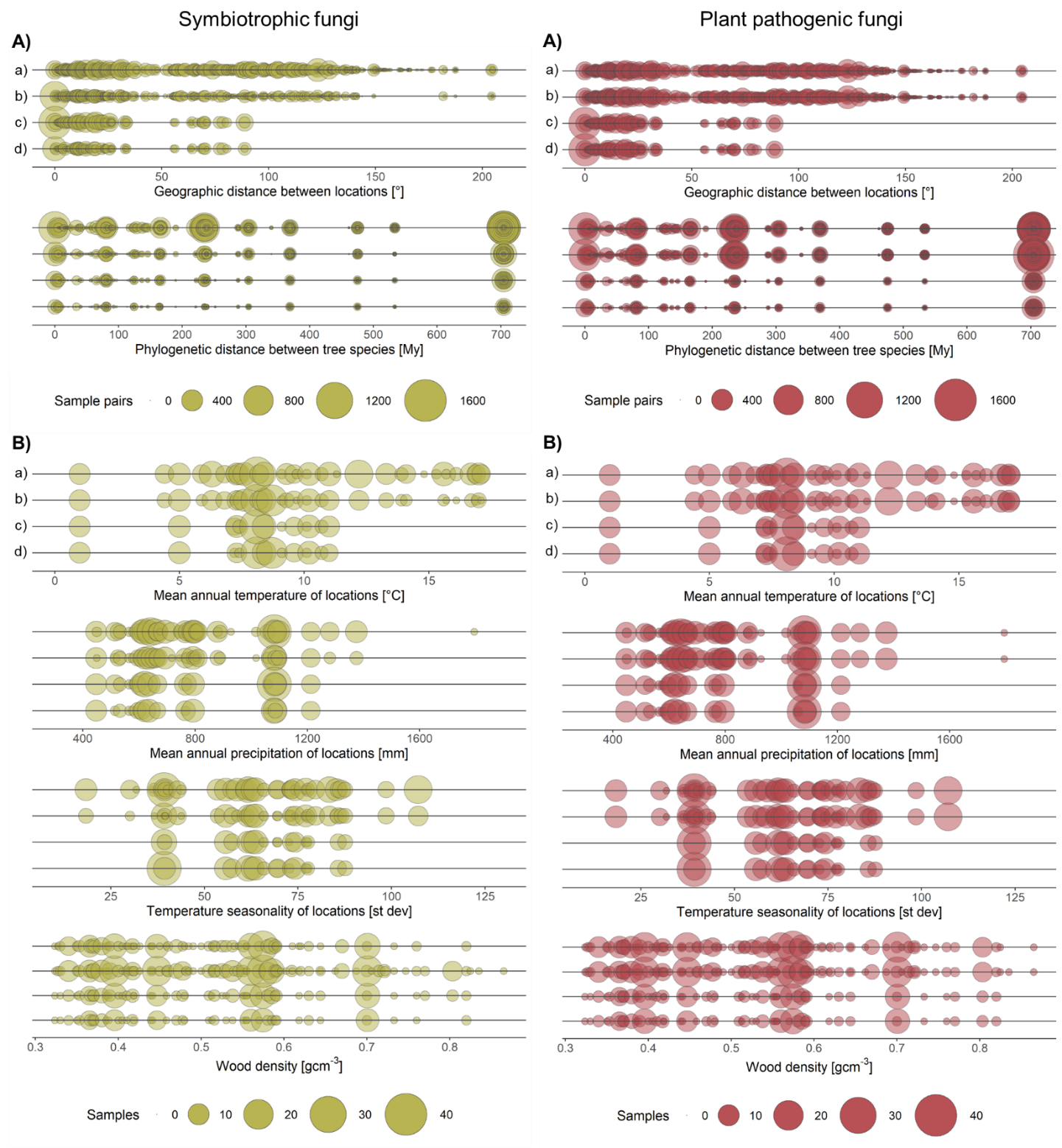

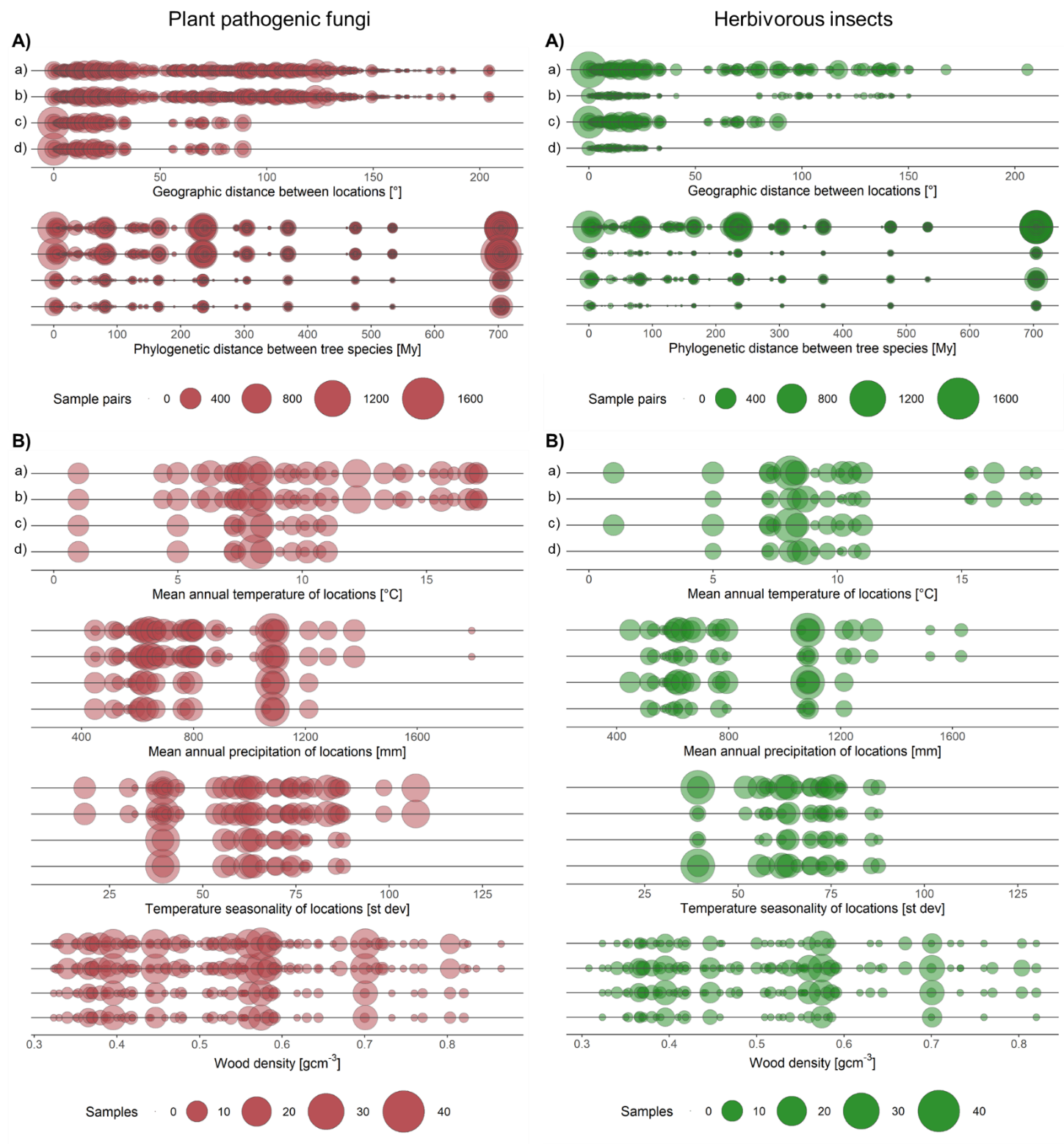
Supplementary Figure S8. **Distribution of collected samples along gradients of different variables. A, B** Frequency of values for pairwise comparisons of host geographic origin (geographic distance) and host species (phylogenetic distance) (**A**) and frequency distribution of climate variables for sites (i.e. mean annual temperature, temperature seasonality and mean annual precipitation) and wood density for tree species (**B**). **a, b, c, d** Distribution of collected samples along gradients of different variables is shown for analysis considering all samples from which organisms were assessed, including blank samples (**a**, “zero adjusted analysis”), all samples from which organisms were assessed and in which they were detected (**b**; “main analysis”), samples from which both insects and fungi were assessed, including blank samples (**c**, “overlap zero adjusted analysis”) and samples from which both insects and fungi were assessed and in which they were detected (**d**, “overlap analysis”). The size of the circles indicates the number of sample pairs (A) or samples (B). Colors indicate different groups (all fungi = grey, saprotrophic fungi = blue, symbiotrophic fungi = yellow, plant pathogenic fungi = red, herbivorous insects = green).

Table S1 **Statistical details on the Generalized Dissimilarity Models (GDMs) in the “main analysis”**. The numbers show model deviance, percent deviance explained by the full model, the p-value of the full model, and the number of permutations used to calculate the statistics for each fitted model (n=50). Numbers are shown for different measures of β-diversity (i.e., Sørensen q=0, Morisita q=1, Morisita-Horn q=2) and species turnover component (i.e., species turnover) and for different functional groups of organisms (i.e., herbivorous insects and all, saprotrophic, symbiotrophic, and plant pathogenic fungi). Numbers in brackets indicate number of samples that were included in the analysis. The percentage of unique variance explained by each variable (i.e., %devexp), the coefficients (i.e., coef; the maximum height of the spline for that variable), the relative value of the coefficients and the p-values (p) are also shown. Bold numbers indicate significant variables.

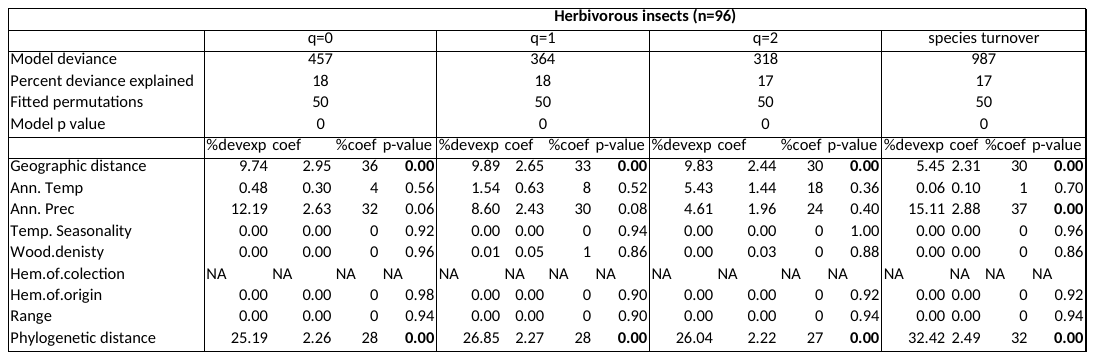

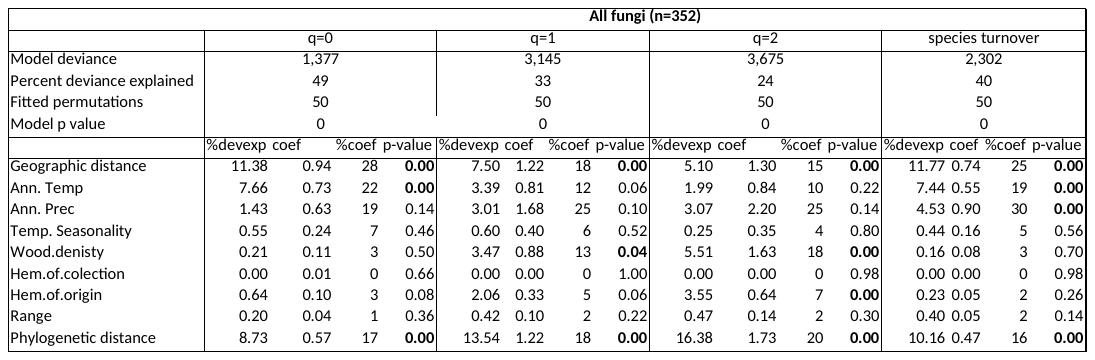

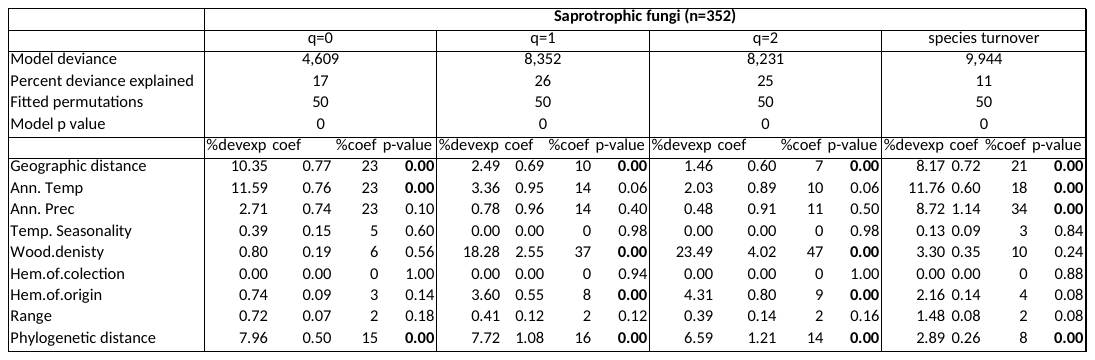

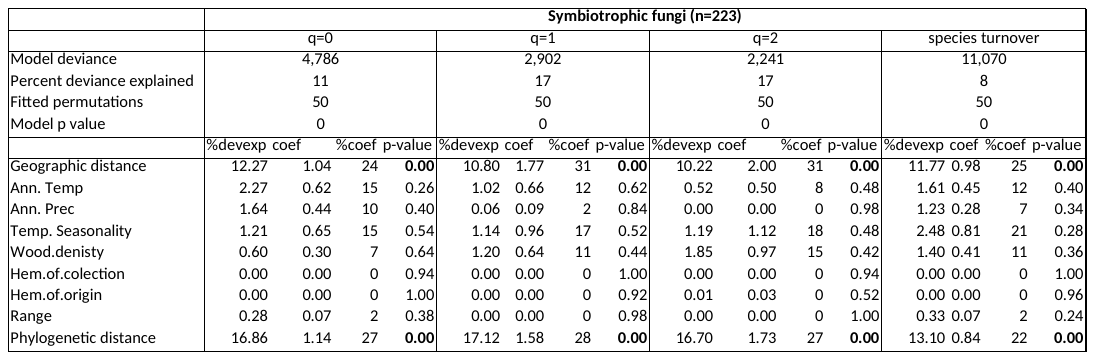

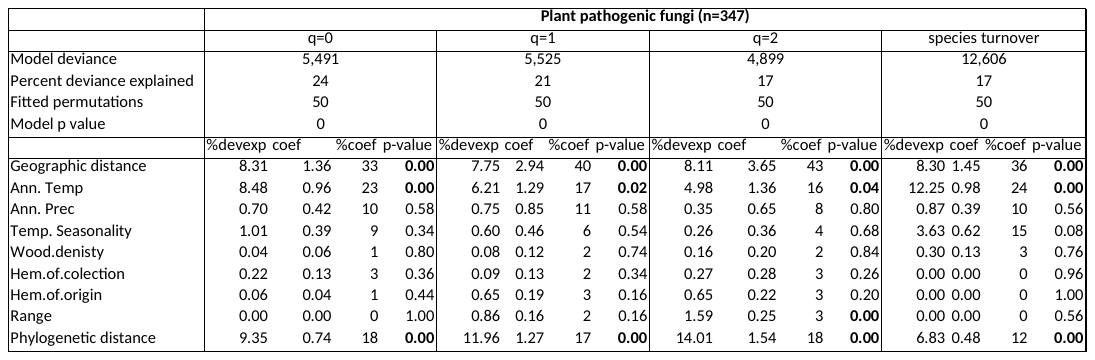

Table S2 **Statistical details on the Generalized Dissimilarity Models (GDMs) in the “zero-adjusted analysis”**. The numbers show model deviance, percent deviance explained by the full model, the p-value of the full model, and the number of permutations used to calculate the statistics for each fitted model (n=50). Numbers are shown for Sørensen (q=0) β-diversity and for different functional groups of organisms (i.e., herbivorous insects and all, saprotrophic, symbiotrophic, and plant pathogenic fungi). Numbers in brackets indicate number of samples that were included in the analysis. The percentage of unique variance explained by each variable (i.e., %devexp), the coefficients (i.e., coef; the maximum height of the spline for that variable), the relative value of the coefficients and the p-values (p) are also shown. Bold numbers indicate significant variables.

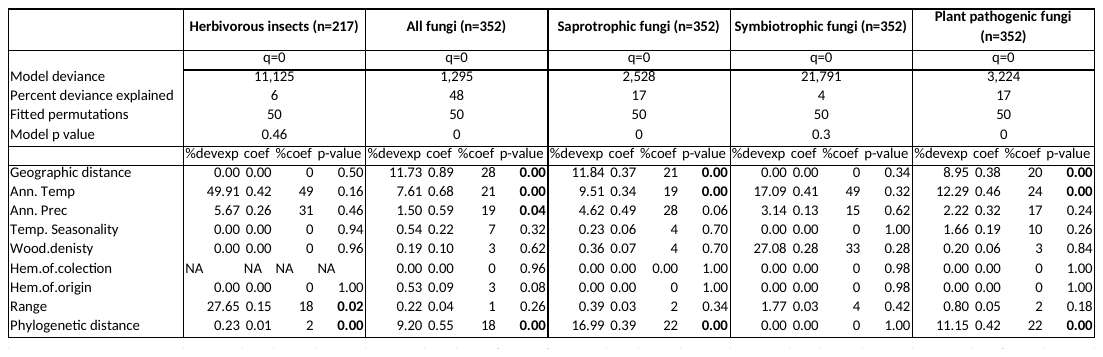

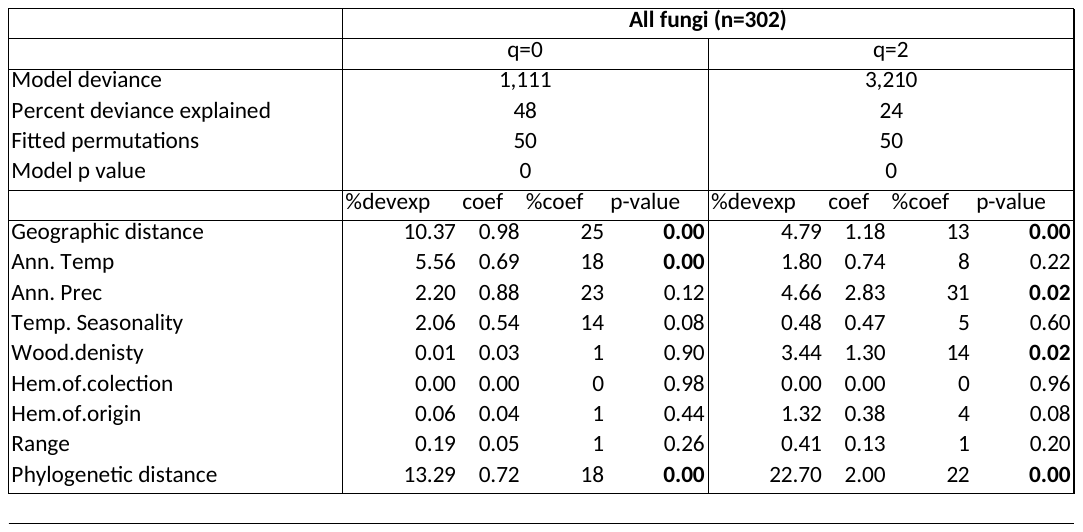
Table S3 **Statistical details on the Generalized Dissimilarity Models (GDMs) in the “rarefied data analysis”**. The numbers show model deviance, percent deviance explained by the full model, the p-value of the full model, and the number of permutations used to calculate the statistics for each fitted model (n=50). Numbers are shown for different measures of β-diversity (i.e., Sørensen q=0 and Morisita-Horn q=2) and for different functional groups of organisms (i.e., all, saprotrophic, symbiotrophic, and plant pathogenic fungi). Numbers in brackets indicate number of samples that were included in the analysis. The percentage of unique variance explained by each variable (i.e., %devexp), the coefficients (i.e., coef; the maximum height of the spline for that variable), the relative value of the coefficients and the p-values (p) are also shown. Bold numbers indicate significant variables.

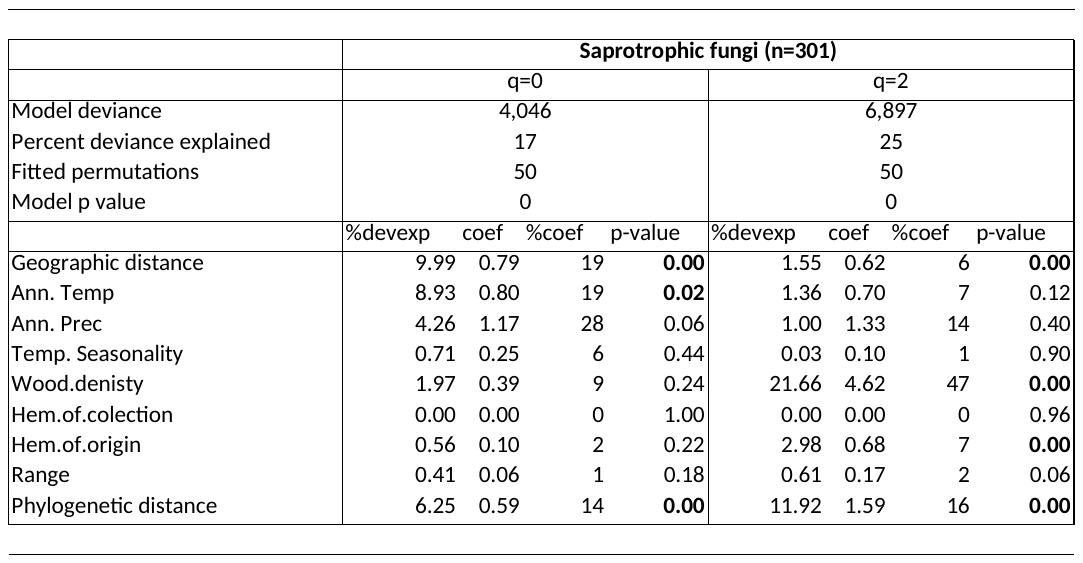

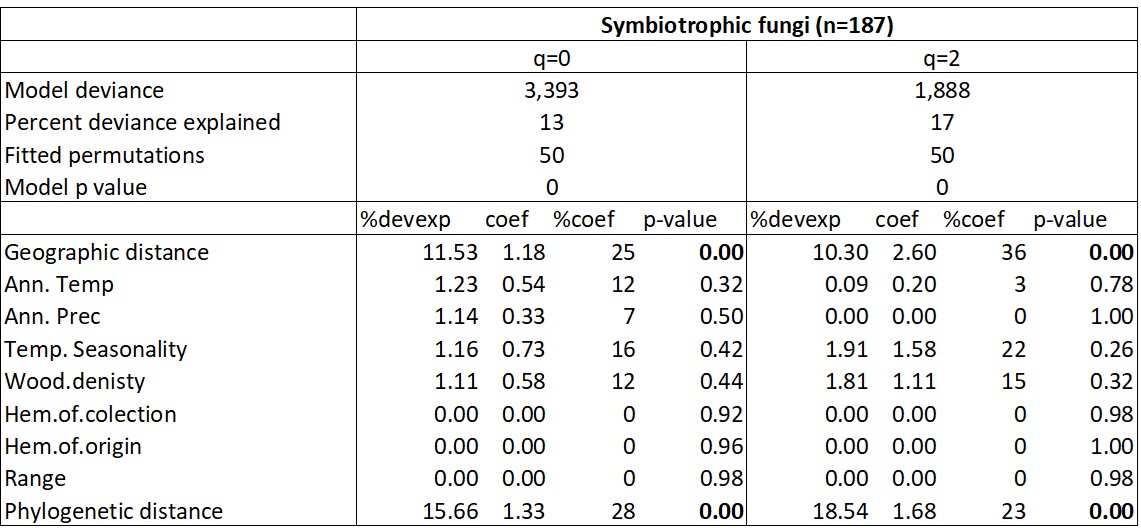

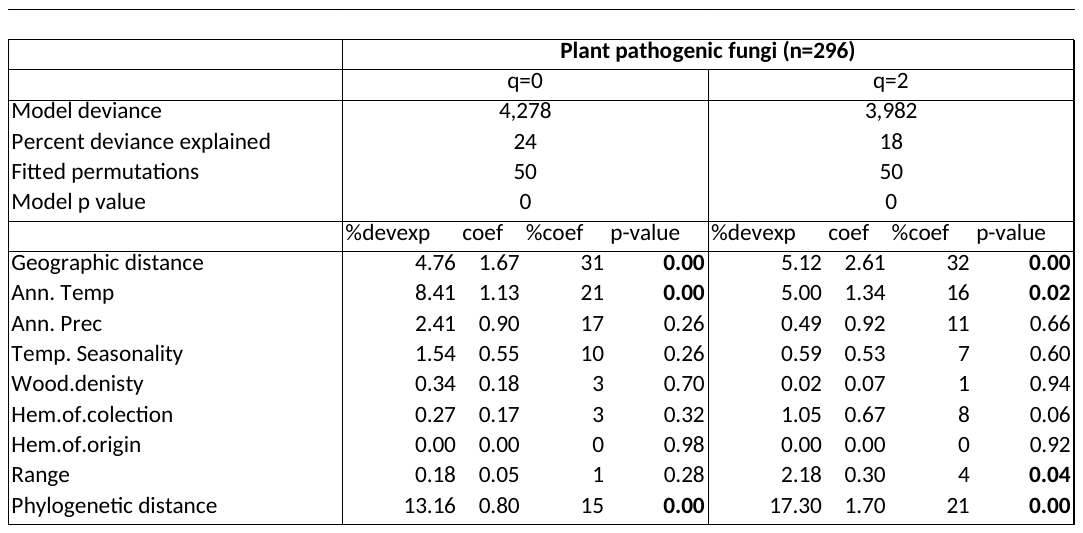

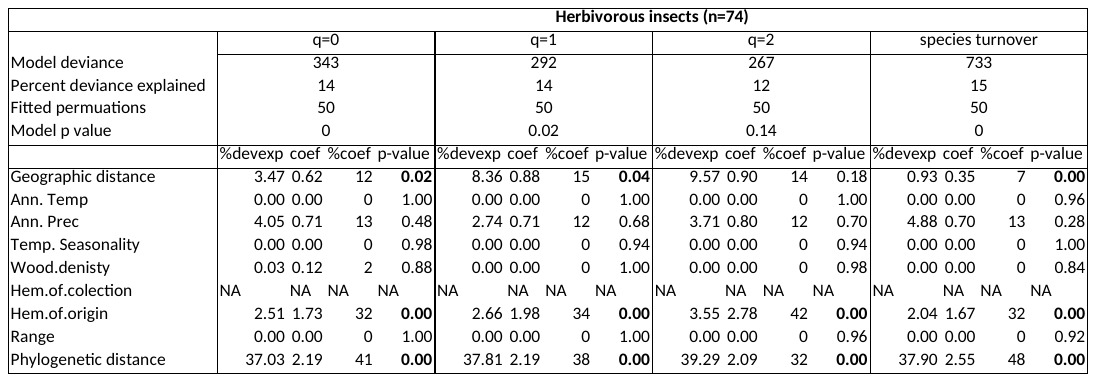
Table S4 **Statistical details on the Generalized Dissimilarity Models (GDMs) in the “overlap analysis”**. The numbers show model deviance, percent deviance explained by the full model, the p-value of the full model, and the number of permutations used to calculate the statistics for each fitted model (n=50). Numbers are shown for different measures of β-diversity (i.e., Sørensen q=0, Morisita q=1, Morisita-Horn q=2) and species turnover component (i.e., betasim) and for different functional groups of organisms (i.e., herbivorous insects and all, saprotrophic, symbiotrophic, and plant pathogenic fungi). Numbers in brackets indicate number of samples that were included in the analysis. The percentage of unique variance explained by each variable (i.e., %devexp), the coefficients (i.e., coef; the maximum height of the spline for that variable), the relative value of the coefficients and the p-values (p) are also shown. Bold numbers indicate significant variables.

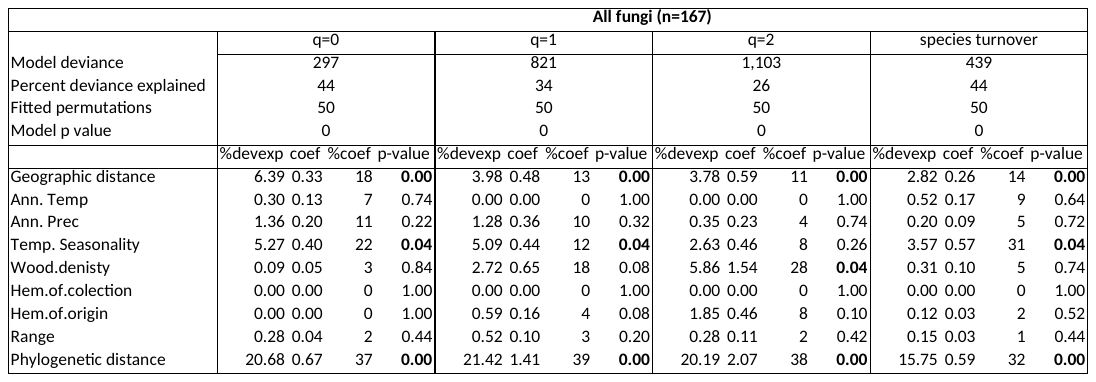

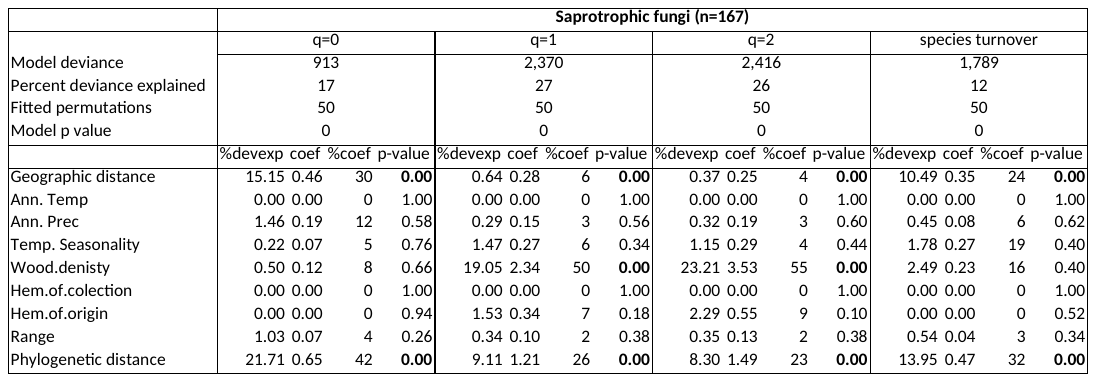

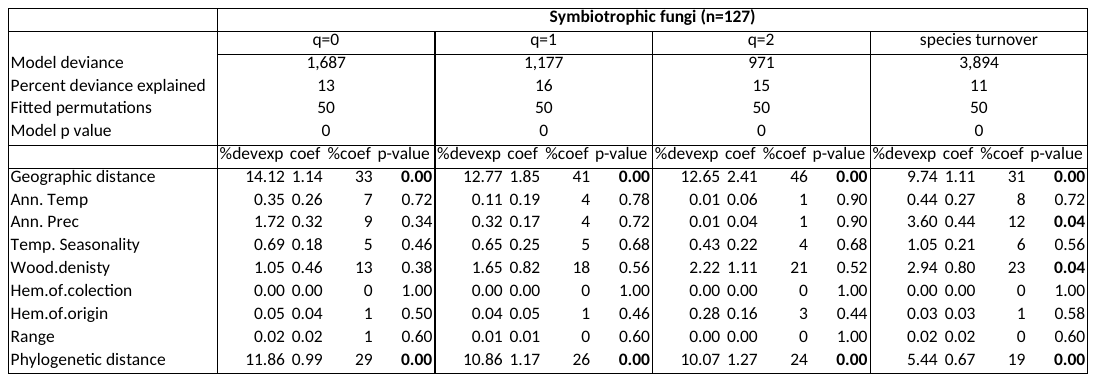

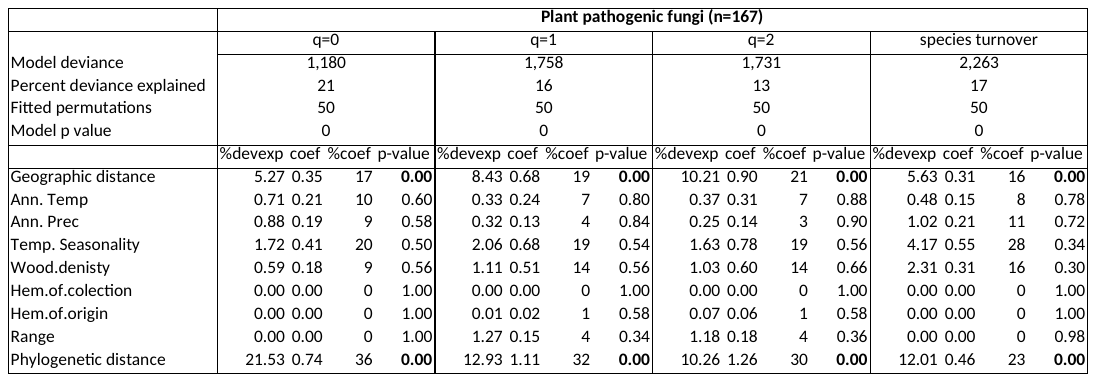

Table S5 **Statistical details on the Generalized Dissimilarity Models (GDMs) in the “zero-adjusted overlap analysis”**. The numbers show model deviance, percent deviance explained by the full model, the p-value of the full model, and the number of permutations used to calculate the statistics for each fitted model (n=50). Numbers are shown for Sørensen (q=0) β-diversity and for different functional groups of organisms (i.e., herbivorous insects and all, saprotrophic, symbiotrophic, and plant pathogenic fungi). Numbers in brackets indicate number of samples that were included in the analysis. The percentage of unique variance explained by each variable (i.e., %devexp), the coefficients (i.e., coef; the maximum height of the spline for that variable), the relative value of the coefficients and the p-values (p) are also shown. Bold numbers indicate significant variables.

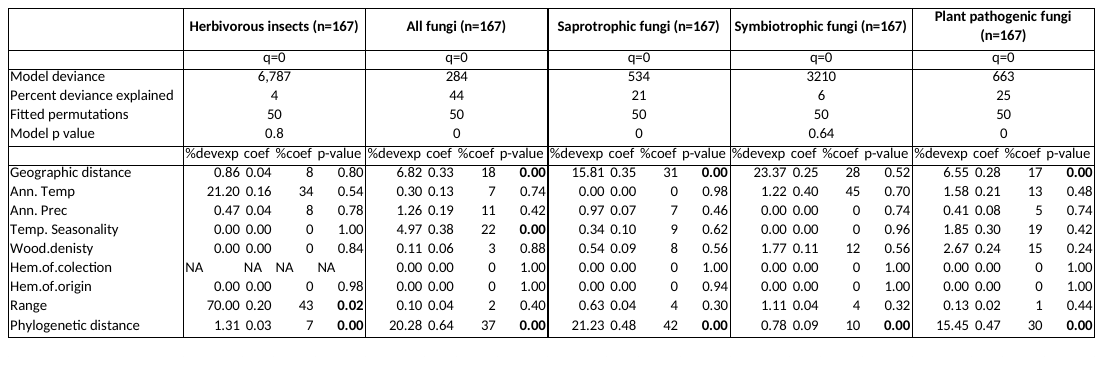
